# Implications of future change scenarios for mosquito-borne disease transmission in the Netherlands

**DOI:** 10.1101/2025.07.09.663836

**Authors:** Marikeni de Wit, Martha Dellar, Gertjan Geerling, Eline Boelee, Mart de Jong, Quirine ten Bosch

## Abstract

**Background:** Mosquito-borne virus transmission is shaped by its ecological context, including land use, climate, and population dynamics. Future changes in these factors may therefore affect the risk and intensity of Usutu virus (USUV) and West Nile virus (WNV) outbreaks in the Netherlands.

**Methodology & findings:** We compared a reference scenario to four future Shared Socio-economic Pathway scenarios developed for the Netherlands, which differed in land use, host distribution, mosquito distribution, and temperature. Temperature (between +1.0°C and +1.7°C) and mosquito abundance (between +5% and +10%) were predicted to increase during the transmission season across the scenarios. Scenario effects on hosts differed between species. We found that outbreak size and growth rate were expected to increase in all future scenarios for both USUV and WNV. These effects were most pronounced early in the season and in scenarios characterised by a high temperature increase and little concern about environmental change. Changes in outbreak risk differed between locations due to spatial variation in changes in host and vector abundance

**Conclusions:** Across a range of possible future scenarios, USUV and WNV outbreaks are expected to become larger, grow faster, and last longer. This is mostly driven by increased temperatures, highlighting the importance of climate mitigation measures to reduce disease outbreak risk and impact.

## Introduction

Transmission dynamics of mosquito-borne diseases are influenced by a complex interplay of ecological factors, including climate, land use, biodiversity, and population dynamics [1]. This ecological context shapes the interactions between the vectors and hosts required for transmission. Therefore, ecological changes can affect the risk of outbreaks and the emergence of new arboviruses [2]. Urbanisation has been associated with increased *Aedes* mosquito density and virus transmission, such as for dengue fever [3]. Similarly, deforestation has been linked to habitat changes which may affect vector and host distributions, between-species contact rates, and movement, and thereby increase outbreak risk [4,5]. One of the most studied environmental drivers of transmission is temperature. Temperature impacts a mosquito’s lifespan, biting frequency, and development, as well as viral replication rates within a mosquito, thereby enhancing transmission potential in multiple ways [6]. These ecological drivers, combined with social drivers like global travel and trade, have contributed to the geographic expansion of mosquito-borne diseases in recent decades [2,7], and are expected to continue to do so [1,8]. Due to the diversity in mosquito and host species and local variation in their distribution, the exact impact of these changes is not straightforward to predict and varies by disease, species, and geographical area.

Two closely related arboviruses that have recently emerged and caused outbreaks in Europe are Usutu virus (USUV) [9] and West Nile virus (WNV) [10]. Both viruses are transmitted in a cycle between mosquitoes and birds, with *Culex pipiens* likely being the most important vector species in Europe [11]. The host ranges of USUV and WNV show a high degree of overlap, with 34 bird species in common across 11 orders [11]. However, while both viruses can infect humans, the impact on human health is larger for WNV as symptoms are more severe [9]. Another important difference is that USUV emergence in Europe has been associated with high Eurasian Blackbird (*Turdus merula*, hereafter: blackbird) mortality [12–15], while such noticeable impacts on bird mortality have not been reported for WNV in Europe. The spread of WNV across Europe has been associated with land use, especially agricultural activity [16], and climate, including spring and summer temperatures and precipitation [17,18].

The Netherlands has seen the emergence of USUV and WNV over the past decade [12,19]. The Netherlands is a water-rich country, with two large rivers forming a delta, with high human and livestock population densities. Such urbanised delta ecosystems are particularly vulnerable to environmental changes, such as increased flooding risk, sea-level rise, and extreme climate events [20]. Dellar et al. [21] developed national One Health scenarios for the Netherlands for 2050 based on the global Shared Socio-economic Pathways (SSPs). SSPs are global scenarios for future societal development, including factors such as demographics, economics, technology, governance, and the environment [22], that can be combined with climate scenarios (Representative Concentration Pathways (RCPs)). The impact of these One Health scenarios has been quantified for land use [23], distribution of several bird species [24], and *Culex pipiens* abundance [25] in the Netherlands. It is, however, not yet known how these changes in temperature and population distributions interact and thereby shape the risk of future USUV and WNV outbreaks. Quantifying the impact of future scenarios on arbovirus disease risk helps in understanding the possible consequences of policy choices and allows for an informed decision-making process. Long-term outbreak predictions can also facilitate preparedness planning.

We leveraged highly comprehensive and detailed One Health scenarios to assess future scenarios of USUV and WNV outbreaks in the Netherlands. We calculated several measures of outbreak risk and intensity and compared these between a reference scenario and four SSPs. These measures were calculated over space and time and stratified by land use class and province.

## Results

### Scenario comparison across season

All future scenarios were characterised by a longer transmission season (up to 17% longer) and higher R_0_ for both pathogens (Figure 1A-B) when compared to the reference scenario. The average R_0_ was highest in SSP3 and lowest in SSP1 for both USUV (reference: 4.0 (95%CI 3.2-4.7), SSP3: 5.2 (95%CI 4.1-6.2), SSP1: 4.7 (95%CI 3.6-5.6)), and WNV (reference: 2.2 (95%CI 1.8-2.9), SSP3: 2.7 (95%CI 2.1-3.7), SSP1: 2.3 (95%CI 1.9-3.1)). Therefore, only results from these two extreme scenarios are presented in the main text (for other scenarios, see Supplementary Material B). R0 surpassed 1 earlier in the season in SSP1 (USUV: 7 days, WNV: 3 days) and even more so in SSP3 (USUV: 16 days, WNV: 9 days). Additionally, R0 also peaks at higher values, most notably for USUV (USUV reference: 11.7 (95%CI 9.4-13.8), SSP3: 13.8 (95%CI 10.9-16.4), SSP1: 12.9 (95%CI 9.9-15.3)) & WNV reference: 6.3 (95%CI 5.2-8.4), SSP3: 7.0 (95%CI 5.6-9.8), SSP1: 6.3 (95%CI 5.1-8.3)). The proportion of locations with R_0_>1 also increased earlier in the season in the future scenarios (supplementary Figure 4C & 9C). Differences in the decline of R0 during September were small, as we assumed that diapause initiation remained constant between scenarios.

**Figure 1:**
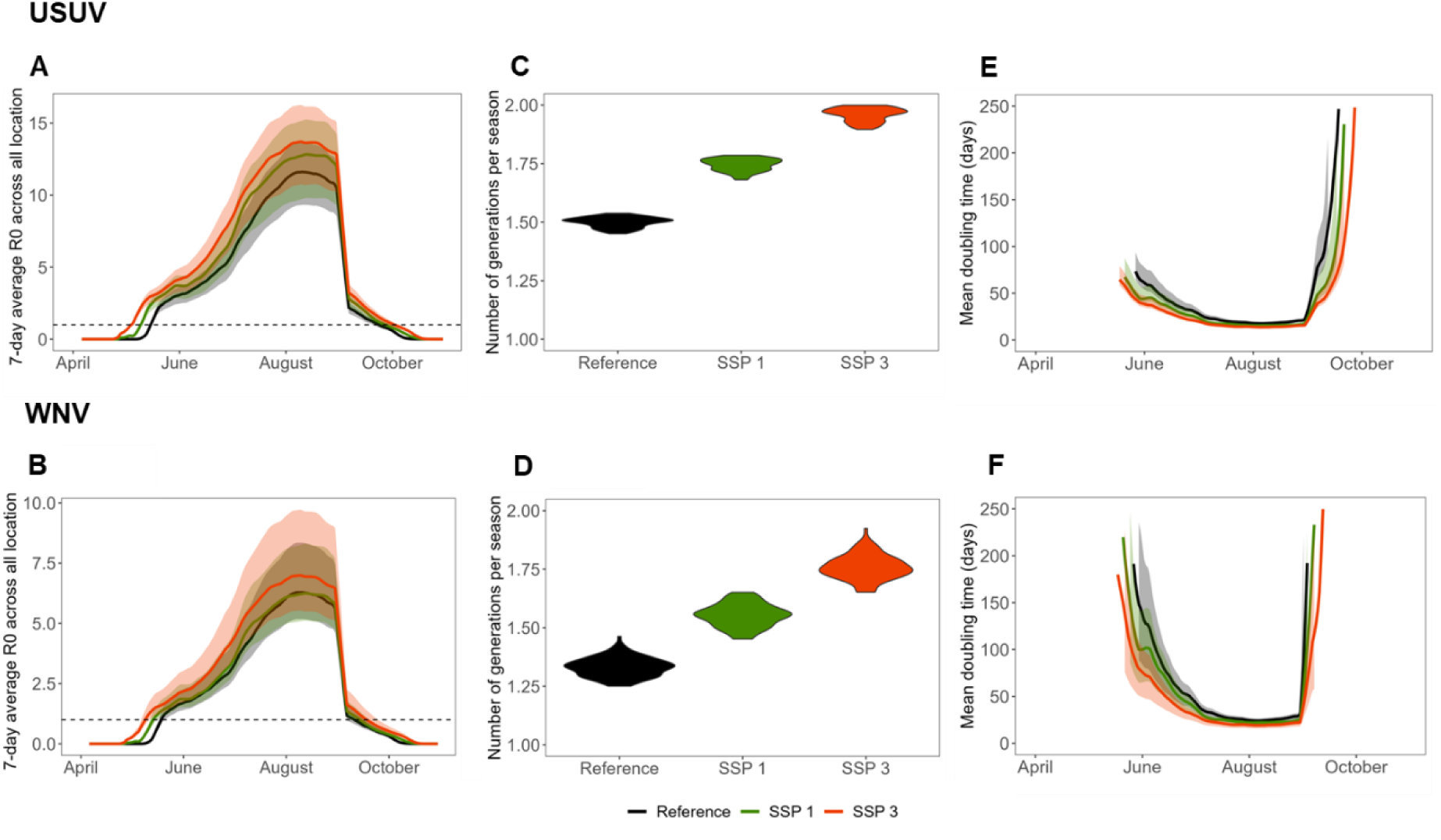
Comparisons of basic reproduction number. (A-B), number of generations per season (C-D), and epidemic doubling time (E-F) between the reference scenario and SSP1 and SSP3. Shaded areas in A, B, E, and F represent the 95%CI based on variation in parameter estimates sampled from posterior distributions. Plots A, C, and E relate to USUV, while B, D, and F relate to WNV.

We estimated a reduction in the generation time across all scenarios (supplementary Figure 4E & 9E), due to higher temperatures. A reduced generation time and increased season length (i.e. period during which R0>1), results in more generations per season. Again, this effect was strongest in SSP3 (Figure 1C-D). The number of generations changed from 1.5 (95%CI 1.5-1.5) in the reference scenario to 1.7 (95%CI 1.7-1.8) in SSP1 and 2.0 (95%CI 1.9-2.0) in SSP3 for USUV and from 1.3 (95%CI 1.3-1.4) in the reference scenario to 1.6 (95%CI 1.5-1.6) in SSP1 and 1.8 (95%CI 1.7-1.9) in SSP3 for WNV. Higher R0 combined with shorter generation time make outbreaks more explosive as the growth rate increases (supplementary Figure 4D & 9D). Especially early in the transmission season, we observed shorter epidemic doubling times. In June, doubling times changed from 41 days (95%CI 28-60) in the reference scenario to 34 (95%CI 24-48) in SSP1 and 28 (95%CI 20-39) in SSP3 for USUV and 75 days (95%CI 43-131) in the reference scenario to 65 (95%CI 39-112) in SSP1 and 47 (95%CI 31-77) in SSP3 for WNV (Figure 1E-F).

To explore the contribution of changes in temperature and mosquito abundance compared to changes in bird abundance, we separated these effects for SSP1 and SSP3. We found that higher temperature and mosquito abundance were the most important drivers of increased R0 (supplementary Figure 6).

### Scenario comparison across space

In the reference scenario, USUV R0 was highest in the South-Eastern region (Figure 2A, provinces of Noord-Brabant (R0 4.8, 95%CI 3.2-5.4), Limburg (R0 4.7, 95%CI 3.9-5.2), and Gelderland (R0 4.4, 95%CI 2.8-5.4)). This is also the area where R0 increased most in both SSP1 and SSP3 for USUV (Figure 2C, supplementary Figure 7). WNV R0 was highest in the South in the reference scenario (Figure 2B, highest value observed in the province of Zeeland (R0 2.6, 95%CI 2.0-2.8)). For WNV, R0 increased most in the region where this was lowest in the reference scenario, which is currently a national park (Figure 2D, supplementary Figure 8). This coincides with the highest increase in (competent) bird abundance (supplementary Figure 1). Whilst WNV R0 increased at the national level, this was not the case for all regions in SSP1. In the Northern and Western regions (provinces of Friesland, Utrecht, Noord-Holland, and Zuid-Holland), R0 remained stable.

**Figure 2:**
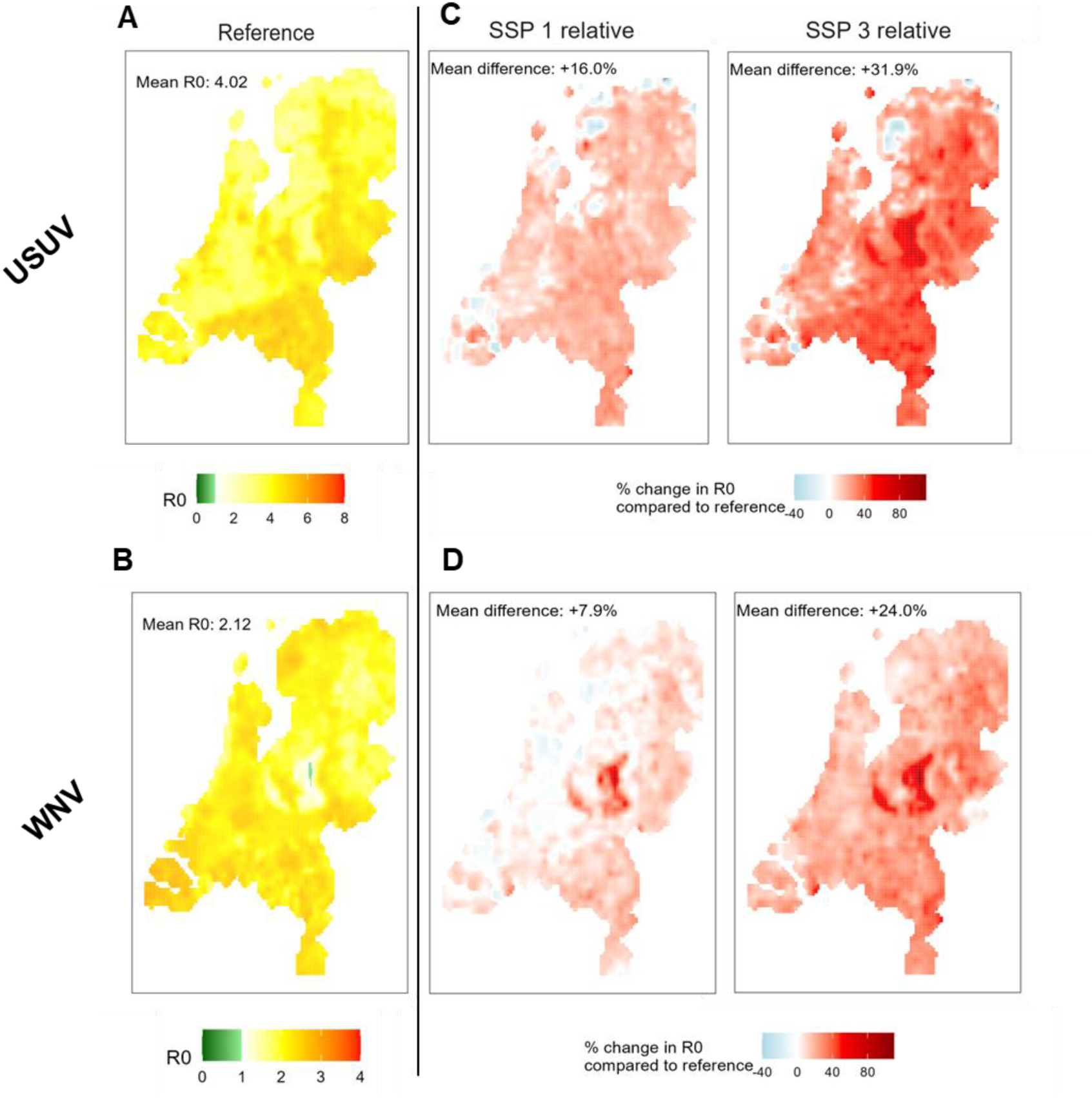
Map of basic reproduction number per grid cell for the reference scenario. (A-B) and the change in the reproduction number between the scenarios and the reference (C-D).

We also compared R0 values between land use classes, which were split into five categories: urban, pasture, crops, forest, and non-forest nature. While differences between land use classes could be observed in bird and mosquito abundance (supplementary Figure 9), differences in R0 values were small (supplementary Figure 10), with non-forest nature and forest being lowest for USUV and WNV respectively. The land use class associated with the highest R0 varied between provinces (supplementary Figure 11, supplementary Figure 12). Relative increases in R0 were similar across land use classes. For WNV, these ranged from 1.04 times in pasture to 1.14 times in forest for SSP1 and from 1.20 times in pasture to 1.33 times in forest for SSP3. For USUV, relative increases ranged from 1.16 times in forest to 1.21 times in non-forest nature for SSP1 and from 1.26 times in pasture to 1.50 times in non-forest nature for SSP3.

### Comparison between USUV and WNV

While R0 increased across all future scenarios for both USUV and WNV, there were local differences between these pathogens. We calculated correlations between these pathogens’ current R0 values and between their predicted change in future scenarios. In the reference scenario, we found low correlation between R0 values for USUV and WNV on a local level (i.e. 5×5km) (Pearson’s r 0.22) and on a provincial level (Pearson’s r 0.29). Correlation between changes in R0 for both pathogens was higher on both a local level (Pearson’s r SSP1 0.53, SSP3 0.73) and a provincial level (Pearson’s r SSP1 0.50, SSP3 0.48).

### Sensitivity analyses WNV host distribution

To assess the implications of our choice of host species for WNV, we included a sensitivity analysis where we assumed mallards and house sparrows were uniformly distributed across the country. Mean R0 values were similar across scenarios (Supplementary Table 1), indicating that assumptions about host distribution do not impact the national-level effect of the scenarios on R0. On a more local level, spatial heterogeneity in the impact of scenarios was naturally reduced (supplementary Figure 13).

## Discussion

Environmental changes can have important consequences for mosquito-borne disease risk. We found that the basic reproduction number increased in all future One Health scenarios developed for the Netherlands for both USUV and WNV, compared to the current situation. Additionally, the duration of the transmission season was projected to be longer, leading to larger outbreaks as the number of generations per season increased. We observed a reduction in the epidemic doubling time, especially early in the season, which makes outbreaks spread more rapidly. The increase in outbreak risk was most pronounced in SSP3 (‘Our town first’), which is characterised by a high increase in temperature and mosquito abundance. Overall increase in outbreak risk was largely driven by increased temperature and mosquito abundance. Spatial variation in risk increase was mostly due to spatial variation in changes in mosquito and bird abundance as temperature changes were more homogeneously distributed across the country. No clear trends were observed between land use classes. There was little correlation in risk between USUV and WNV. The sensitivity analysis regarding the choice (and therefore distribution) of host populations for WNV showed that while local estimates were sensitive to these distributions, national trends of R0 between scenarios were robust to these.

Experimental studies suggest that WNV transmission will peak at average daily temperatures around 24.5°C [26,27]. As mean daily temperatures of 24.5°C were not reached even in SSP3, transmission was projected to increase in all scenarios and continued temperature rise is likely to further increase virus transmission. A European-wide study that modelled WNV risk for different climate change scenarios found an up to 3.5 times increase in population at risk of WNV infection in 2040-2060 compared to 2000-2020, with the largest risk increase observed in Western Europe [28]. Similar to our study, they observed the smallest increased risk for the low emissions scenario (RCP 2.6) and the largest increase in the high emissions scenario (RCP 8.5). However, this study did not incorporate land use changes and future host and vector abundance estimates. Secondly, this study focused on WNV geographical expansion, using a binary indicator of infection risk, instead of a continuous value such as R_0_.

We did not observe clear differences in transmission risk between land use classes. Land use explained about 37% of variation in abundance in all three bird species [24] and almost 50% of variation in *Cx pipiens* abundance [25]. However, while species abundance differed between land use classes, the effect of land use varied between species. Their combined association with transmission risk was therefore reduced. This effect may be different in regions with different dominant host species. The choice of grid size may also affect the lack of observed variation in transmission risk between land use classes, as adult mosquito abundance has been found to vary locally between park and residential spaces within urban regions [29].

Many studies aimed at projecting future outbreaks have not been validated on historic or current surveillance data. For USUV, we used a validated model to derive parameter values used in the Next Generation Matrix, thereby improving the reliability of our results. Additionally, we built on previously published projections of mosquito and bird abundance, which were not only based on temperature scenarios, but also changes in land use and other environmental variables. The embedding of our scenarios into the information-rich SSP scenarios specifically developed for the Netherlands, allows for a contextualization of results beyond quantitative indicators. However, the exact values of R0 and R0-based indicators should be interpreted with care, especially for WNV. While parameter estimates for USUV were obtained from an USUV transmission model calibrated to the Dutch context [30], such calibrated parameters were not available for WNV, due to the currently limited transmission in the Netherlands. We therefore relied on published literature for infection parameters and assumed a similarly sized competent host population as compared to USUV. Additionally, a wide range of bird species have been found competent for USUV and WNV transmission. These species differ in their competence and distribution, thereby leading to spatial heterogeneity in transmission risk. Our sensitivity analysis showed that the choice of host species affected the spatial pattern in disease risk, yet not the general trends. In reality, these spatial patterns are not only affected by distribution of competent hosts, but also of non-competent hosts. Due to a lack of information on the distribution of all possible *Cx pipiens* hosts, we assumed the total host population to be the same in each location. We also assumed that future changes in the competent bird species were reflective of changes in the total host population and therefore, in future scenarios, kept the overall ratio of competent vs non-competent hosts equal to the reference scenario. While these assumptions are unlikely to impact the overall differences between scenarios, they do impact local-level projections. It is also important to note that results are based on the basic reproduction number and therefore the metrics presented do not account for immunity in the population. Given the ongoing circulation of both viruses in the Netherlands [31], immunity will play an important role in shaping future transmission. Other possible directions for future extensions include exploring the impact of future scenarios on diapause initiation and duration as well as on bird movement patterns. Moreover, we compared scenarios based on the mean generation interval, but more detailed approaches for calculating the distribution of generation intervals, such as presented in [32,33], could be adapted to USUV and WNV to study this in greater detail.

Our results indicate that WNV and USUV outbreaks are expected to spread more rapidly in the future. This means that timely surveillance becomes more challenging and the window of opportunity for response becomes smaller. Should human vaccines become available in the future, these will have to be rolled out faster if a reactive vaccination strategy is followed. The observed increase in transmission between mosquitoes and birds is likely to also lead to more spillover cases to other animals, including humans. Future scenarios do not only differ in their projected transmission risk, but also in their ability to anticipate and respond to outbreaks [21]. The smallest increase in transmission risk was observed in SSP1. Health services in this scenario are also well-prepared for outbreaks and have effective early-warning systems [34]. People are generally healthy and adhere to public health messages. This contrasts with SSP3 where transmission risk increased the most. The impact of larger outbreaks in this scenario is further exacerbated by a lack of a coordinated response, a less healthy population, and lower adherence to public health messages. The predicted increase in future transmission risk is likely to also apply to other arboviruses. For example, optimal temperature for transmission of Rift Valley Fever is around 26°C [27], which is above current mean daily temperatures. Higher temperatures also increase the suitability for exotic mosquito species in Europe, including *Ae aegypti* and *Ae albopictus* [8,35], known vectors for dengue and chikungunya viruses. Combined with high levels of travel and trade, as is predicted especially for SSP4 and SSP5, this leads to a higher risk of newly emerging arboviruses.

### Conclusion

Across all future scenarios, USUV and WNV outbreaks are expected to become larger, last longer, and spread more rapidly, especially in the southern and eastern parts of the Netherlands. This is mostly driven by increased temperatures, suggesting that climate mitigation measures are an effective strategy to reduce outbreak risk and impact. These findings highlight the need for increased preparedness for arbovirus outbreaks in the future. The development and implementation of surveillance and response activities, including potential vaccines or treatment options, is important to mitigate future arbovirus risks for both humans and animals.

## Methods

Local transmission is governed by interactions between hosts and mosquitoes and is sensitive to temperature. Therefore, host abundance, mosquito abundance, and temperature were obtained for the different scenarios for the Netherlands. To calculate local differences in outbreak risk measures, the country was divided into 5×5 km grid cells. As abundance and temperature change over time, we calculated all measures over space and time. Outbreak risk measures were calculated daily for each grid cell in the Netherlands from April to November. The reference scenario represents the current situation, while the future scenarios represent 2050. Scenario impacts were quantified using several key epidemiological metrics. Local outbreak risk was determined by calculating the basic reproduction number (R_0_), which determines the average number of secondary cases resulting from one average infectious individual in a fully susceptible population. The explosiveness of an outbreak was quantified by calculating the epidemic doubling time (i.e. the time it takes for the outbreak to double in size in a fully susceptible population).

## Model overview

We developed a mathematical, compartmental model of USUV and WNV transmission between *Culex pipiens* mosquitoes and bird host populations. Dead-end hosts, such as humans and horses, were not explicitly included in the model as they do not contribute to onwards transmission. In this model, virus transmission occurred within grid cells, assuming a homogeneous distribution of birds and mosquitoes within these cells. Birds can get infected when they get bitten by an infectious mosquito, while mosquitoes can get infected when they bite an infectious bird. We did not include connections between cells. The mosquito mortality rate, extrinsic incubation period, and biting rate were assumed to be temperature dependent. Parameter values were obtained from literature for both USUV and WNV (Table 1, Table 2). R0 was calculated from this model for each location and day using the Next Generation Matrix (NGM) approach (Supplementary Material A). Each element in the NGM, *k_ij_*, represents the expected number of infections of type *i* caused by an individual of type *j*. The dominant eigenvalue of this matrix equals R0.

**Table 1:** Parameters and their values used in the NGM for USUV. T indicates temperature, t indicates time. HDI = highest density interval.

| Parameter | Description | Value | Source |
| --- | --- | --- | --- |
| **Blackbirds** |  |  |  |
| $\delta_b(t)$ | Daily number of births | 1 May – 15 June:<br>0.056 per bird<br>Else: 0 | Value: see supplementary material.<br>Timing: [38] |
| $\mu_{JB}$ | Mortality rate, juvenile | 0.0059 / day | [38] |
| $\mu_{AB}$ | Mortality rate, adult | 0.0011 / day | [39] |
| $\rho_{bm}$ | Transmission probability per bite, mosquito to blackbird | 0.88 | [40] |
| $\varepsilon$ | Intrinsic incubation rate | 0.67 / day | [41,42] |
| $\gamma$ | Recovery rate | 0.25 / day | [43] |
| $v_b$ | Disease-induced mortality rate | 0.79 / day<br>(95% HDI 0.60-1.0) | Posterior distribution obtained from [30] |
| **Additional reservoir population** |  |  |  |
| $\delta_r(t)$ | Daily number of births | 1 May – 15 June:<br>Variable, depending on value for mortality<br>Else: 0 | Value: see supplementary material.<br>Timing: [38] |
| $\mu_{JR}$ | Mortality rate, juvenile | 0.0059 / day | Set equal to blackbird juvenile |
| $\mu_{AR}$ | Mortality rate, adult | $4.0 \times 10^{-4}$ / day<br>(95% HDI $1.5 \times 10^{-3}$ – $2.2 \times 10^{-4}$) | Posterior distribution obtained from [30] |
| $\rho_{bm}$ | Transmission probability per bite, mosquito to reservoir host | 0.88 | [40] |
| $\varepsilon$ | Intrinsic incubation rate | 0.67 / day | [41,42] |
| $\gamma$ | Recovery rate | 0.25 / day | [43] |
| $v_r$ | Disease-induced mortality rate | 0 | Posterior distribution obtained from [30] |
| $\omega$ | Relative biting frequency | 34.8<br>(95% HDI 27.0-45.2) | Posterior distribution obtained from [30] |
| **Culex pipiens** |  |  |  |
| $\mu_m(T)$ | Mortality rate [daily] | For $T < 33$ °C:<br>$1 / (-4.86 \cdot T + 169.8)$,<br>else 0.17 | [27] fitted to data from [44–46] |
| $b(T)$ | Biting rate [daily] | $0.344 / (1 + 1.231 \cdot \exp(-0.184 \cdot (T - 20)))$ | [42] fitted to data from [47] |
| $q$ | Proportion of bites on competent host | Variable | Calculated, see supplementary material. |
| $\rho_{mb}$ | Transmission probability per bite, blackbird & reservoir population to mosquito | 0.82<br>(95% HDI 0.45-1.0) | Posterior distribution obtained from [30] |
| $u(T)$ | Extrinsic incubation rate [daily] | For $T > 13$ °C:<br>$7.38 \times 10^{-5} \cdot T \cdot (T - 11.4)(45.2 - T)^{1/2}$,<br>else 0 | [27] fitted to data from [48,49] |
| $\sigma(t)$ | Rate of entering diapause | After 1 September:<br>0.05. Else: 0 | [50] |
| $\alpha$ | Abundance scaling<br>parameter | 0.11<br>(95%HDI 0.06-0.15) | Posterior distribution<br>obtained from [30] |

**Table 2:**
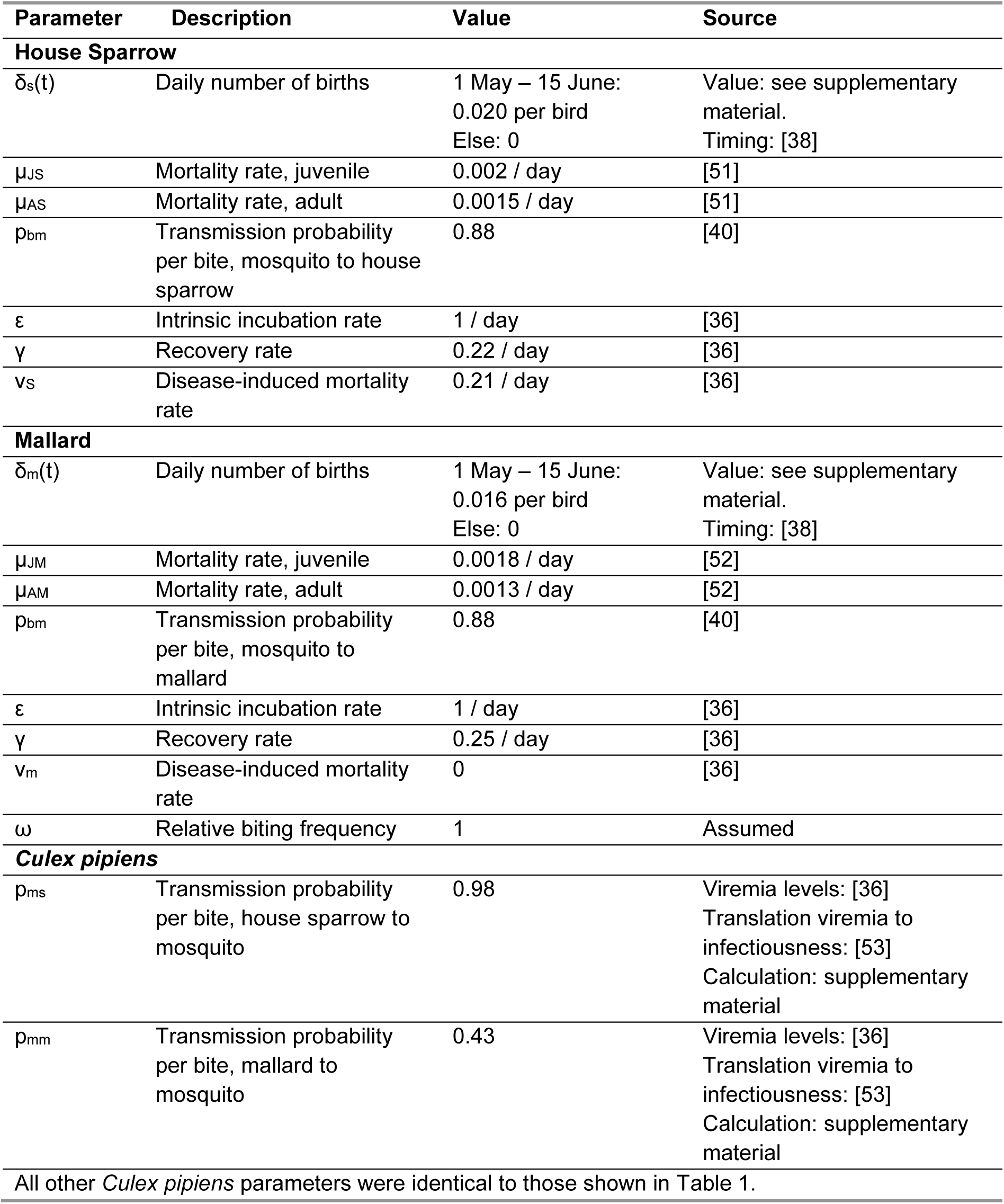
Parameters and their values used in the NGM for WNV. T indicates temperature, t indicates time. HDI = highest density interval.

The host population for USUV consisted of two groups: blackbirds and an additional reservoir population. Blackbirds were included as a host species because high prevalence and high USUV-related mortality has been observed in this population [12,13,31]. An additional reservoir population was included, because it has been shown that this population played a large role in USUV transmission in earlier outbreaks in the Netherlands [30]. The host population for WNV also consisted of two host groups which differed with respect to their infectiousness and mortality rate. The ‘high infectiousness’ population was represented by House Sparrows (*Passer domesticus*, hereafter: house sparrow), while the ‘low infectiousness’ population was represented by Mallards (*Anas platyrhynchos*, hereafter: mallard) [36]. Mortality rate due to WNV infection was higher in house sparrows than in mallards [36]. Further information on this choice of species for WNV is presented in Supplement A. Due to differences in survival rates across life stages, we divided all bird populations into juveniles (from fledging until the next breeding season) and adults (birds older than one year).

*Culex pipiens* mosquitoes bite a wide range of hosts [37], including hosts unable to further transmit the virus (i.e. non-competent hosts). To account for the presence of non-competent hosts, we calculated the proportion of mosquito bites taken on competent hosts. We assumed that the modelled bird populations represent all competent hosts and that the total number of (competent and non-competent) hosts is the same in each location. We assumed that the proportion of bites on competent hosts is proportional to the relative abundance of competent host species in a grid. To reflect this, we calculated this proportion as the relative abundance compared to the cell where abundance was highest, with the grid cell with the highest competent host abundance set to 100%.

### Development of future change scenarios

We compared USUV and WNV transmission potential, over time and space during the transmission season, under a reference scenario and four future scenarios for the Netherlands. The reference scenario represents the current situation, while the future scenarios are based on the Dutch One Health scenarios derived from the global Shared Socio-economic Pathways (SSP) by Dellar et al. [21]. The scenarios were developed by combining existing scenarios (including global and European SSPs), planned (inter)national policies, (grey) literature, and stakeholder consultations. A total of four scenarios were developed: SSP 1 (“Together green”), SSP 3 (“Our town first”), SSP 4 (“The green gulf”), SSP5 (“After us comes the deluge”) (Figure 3).

**Figure 3:**
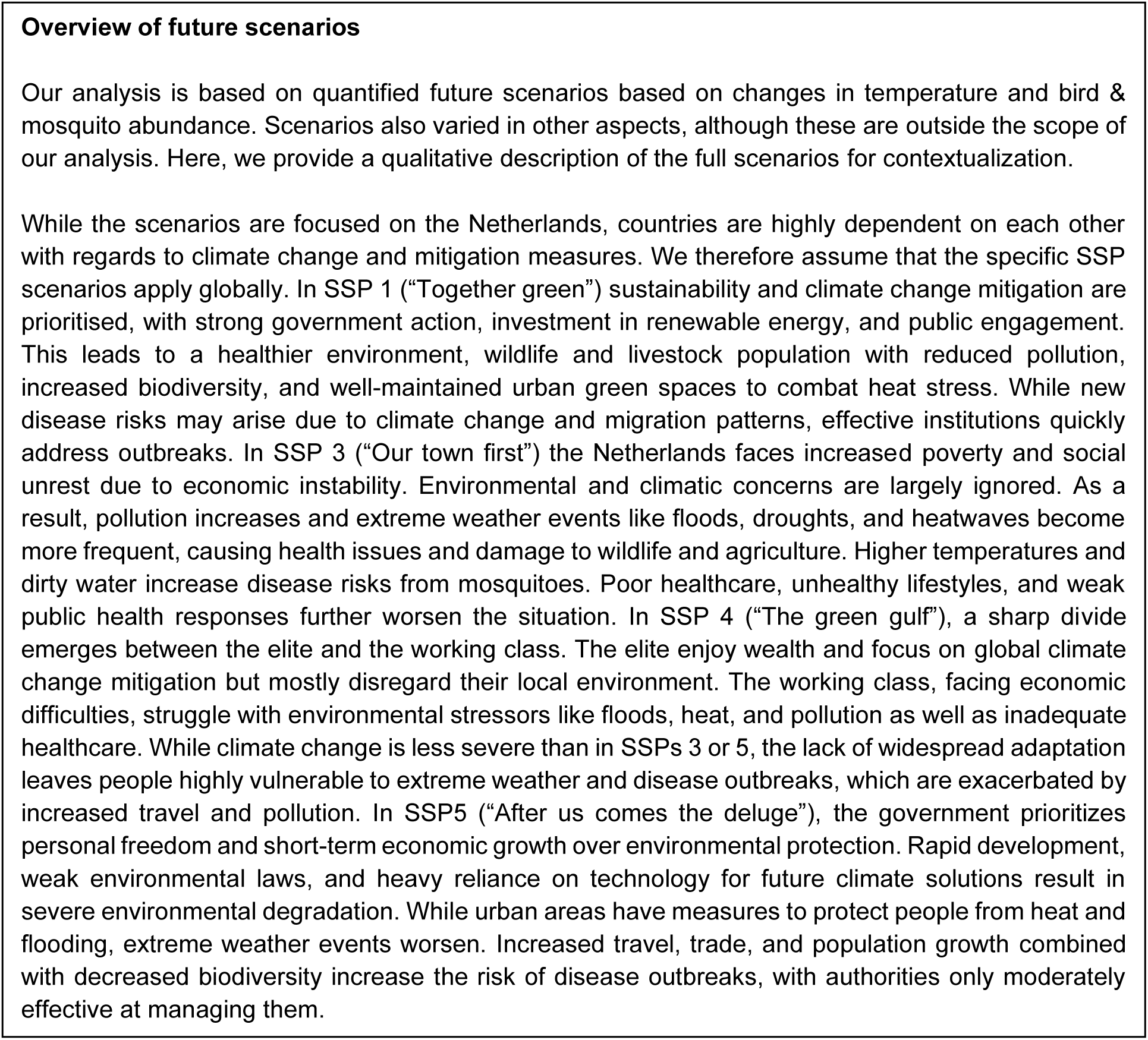
Qualitative description of future One Health change scenarios for the Netherlands. A more detailed description is available in Dellar et al. [21].

These SSP scenarios were used to derive future mosquito and bird abundance and distribution. Impacts of scenarios were quantified based on changes in a wide range of factors, including human population, vegetative cover, water quality, and climate. These changes were derived from scenario-specific land use maps [23], qualitative SSP scenarios [21] and climate scenarios from the Royal Netherlands Meteorological Institute (KNMI) [54]. Scenario-specific blackbird, house sparrow, mallard, and *Cx. pipiens* abundance maps were developed that incorporate these changes ([24] for birds, [25] for *Cx. pipiens*). Bird distributions were created for the breeding season using Random Forest models fitted to point count data from the Netherlands (Meetnet Urbane Soorten [55], Meetnet Agrarische Soorten [56] and Common Bird Census [57] from Sovon Vogelonderzoek Nederland (Dutch Centre for Field Ornithology)) and France (Common Bird Monitoring Scheme [58]), using a set of environmental and climatic predictors. Data from France were included because the current French climate is deemed similar to the future Dutch climate. Mosquito distributions were created using a similar approach where data on female mosquito trap counts was used from the National Mosquito Survey 2010-2013 [59] and the MODIRISK project [60].

Future climate data was taken from scenarios developed by the Dutch Meteorological Organisation for the period 2036-2065 [54]. This consists of six scenarios: low, medium, and high greenhouse gas emissions, each with a wet and dry variant. The different emissions levels are approximately equivalent to RCPs (Representative Concentration Pathways) 2.6, 4.5 and 8.5 respectively. Each scenario is the result of an 8-model ensemble and uses 1991-2020 as a reference period. The scenarios are on a 12 km grid, which we converted to a 5 km grid using bilinear interpolation. For our analysis, SSP 1 was paired with the low emissions scenarios, SSP 4 was paired with the medium emissions scenario, and SSPs 3 & 5 were paired with the high emissions scenarios. To calculate temperature-dependent parameters, we averaged over the 30 years, the 8 ensembles and the wet and dry variants. Future scenarios thereby represent the average predicted temperature in 2050. The reference scenario was defined as the average across the KNMI reference period of 1991-2020.

Apart from the temperature-dependent parameters, parameter values remained the same across scenarios. Initiation and end of diapause was also kept equal to the reference scenario, because these processes, at least partially, depend on day length which does not change in future [61]. Due to a lack of information on the future distribution of the non-competent host population, the proportion of bites on competent hosts averaged across all grid cells was also kept equal to the reference scenario. Locally, the proportion of bites on competent hosts could still differ between scenarios.

### Scenario-specific abundance and parameters

Abundance, temperature, and therefore all temperature-dependent parameters, differed between scenarios. Overall, blackbird abundance remained relatively similar across all scenarios, while both mallard and house sparrow abundance increased (Figure 1A-C). The largest increase compared to the reference scenario was observed in SSP1 for house sparrows. On a local level, house sparrow and mallard abundance increased almost everywhere, while the limited changes in blackbird abundance were a combination of local increases and decreases (Supplementary Figure 1). Across all scenarios, mosquito abundance increased between 4.7% (95%CI 3.3-6.1) in SSP1 and 9.6% (95%CI 8.3-11.0) in SSP3. This increase was most prominent early in the year, while peak abundance was somewhat reduced compared to the reference scenario (Figure 1D, spatial results: supplementary Figure 2). Temperature-dependent parameters were highest for SSP3 and 5 as these scenarios are associated with the highest temperatures (Figure 1E). Average temperature increases across the period April-November ranged from +1.0°C (95%CI 1.00-1.05) in SSP1 to +1.7°C (95%CI 1.66-1.71) in SSP3 and 5 (supplementary Figure 3).

### Analyses

Scenarios were compared based on three important metrics of potential pathogen transmission: the basic reproduction number *R_0_*, the generation time *T*, and the epidemic growth rate *r*.

#### Basic reproduction number

Basic reproduction numbers (R_0_) were calculated using the Next Generation Matrix approach [62] (supplementary material A). R_0_ indicates the average number of secondary cases resulting from one average infectious individual in a fully susceptible population. Uncertainty in parameter estimates was incorporated in R_0_ estimates by sampling 100 parameter sets from the posterior distributions (Table 1) taken from [30].

#### Generation time

The generation time *(T*) is defined as the time interval between successive infections in hosts. We derived the mean generation time for both viruses as the sum of four sequential steps in the transmission cycle, adapted from [32].

- Intrinsic incubation period This period was defined as the time between a host getting infected and becoming infectious. This was calculated by taking the inverse of the intrinsic incubation rate (ε) (Table 1 & Table 2). When estimates varied between host species, we used the average.
- Host-to-mosquito transmission period This period was defined as the time between a host becoming infectious and a susceptible mosquito getting infected from biting this infectious host. Assuming a constant infectiousness and hosts being bitten at a constant rate during the infectious period, a mosquito infection occurs, on average, halfway through the infectious period. We therefore defined the mean host-to-mosquito transmission period as half of the host infectious period. The host infectious period was calculated as the sum of the inverse of the recovery rate (γ), disease-induced mortality rate (ν), and natural mortality rate (μ) (Table 1 & Table 2). Again, when estimates varied between host species, we used the average.
- Extrinsic incubation period This period represents the time between a mosquito getting infected and becoming infectious. This was calculated by taking the inverse of the extrinsic incubation rate (u(T)) (Table 1 & Table 2).
- Mosquito-to-host transmission period This period was defined as the time between a mosquito becoming infectious and transmitting the infection to a susceptible host. When we assume a constant infectiousness and mortality rate, transmission occurs, on average, halfway through the infectious period (i.e. halfway through a mosquito’s lifespan). We therefore defined the mean mosquito-to-host transmission period as half a mosquito’s lifespan, calculated by taking the inverse of the mortality rate (μ_m_(T)) (Table 1).

#### Epidemic growth rate

While the reproduction number provides information on the number of secondary cases arising from an infectious individual, it does not provide information about how quickly these secondary cases occur. The epidemic growth rate is defined as the rate of change in the (log-transformed) number of new cases per unit of time and provides insight into the explosiveness of an epidemic. The relationship between the epidemic growth rate and R_0_ depends on the distribution of the generation interval [63]. Assuming that all generation intervals are equal to the mean generation interval, the relationship between R_0_ and the growth rate can be expressed as

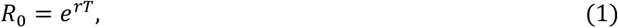

where *r* is the epidemic growth rate, and *T* is the mean generation interval. The growth rate is directly related to, the more intuitive, epidemic doubling time which represents the time it takes for the number of cases to double. This can be expressed as

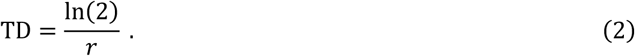

### Sensitivity analyses

We conducted a sensitivity analysis to assess the implications of our choice of host species for WNV. Here, we assumed the host population to be uniformly distributed across the country. The average proportion of bites on competent hosts was kept equal to the other scenarios.

## Author contributions

**MdW**: conceptualisation, methodology, software, formal analysis, visualisation, data curation, writing – original draft, writing – reviewing and editing. **MD**: conceptualization, data curation, writing – reviewing and editing. **GG:** conceptualisation, writing – reviewing and editing. **EB**: conceptualisation, writing – reviewing and editing **MdJ**: conceptualisation, methodology, supervision, writing – reviewing and editing. **QtB**: conceptualisation, methodology, supervision, writing – reviewing and editing.

## Data availability statement

All data used in this study have been published before. Future climate data was taken from scenarios developed by the Dutch Meteorological Organisation for the period 2036-2065 (van Dorland et al., KNMI National Climate Scenarios 2023 for the Netherlands. De Bilt: 2024.). Data for these scenarios, including the reference scenario, is available from the KNMI website (https://klimaatscenarios-data.knmi.nl/downloads). *Cx pipiens* abundance estimates have been published in Krol et al. 2024 (https://doi.org/10.21203/RS.3.RS-5298493/V1) and are available upon request from the corresponding author. Bird abundance estimates have been published in Dellar et al. 2024 (https://doi.org/10.1007/s10393-025-01727-9). For this analysis, all model inputs (with the exception of the bird data), model outputs, code and supplementary information can be accessed via the Dryad repository: https://doi.org/10.5061/dryad.r2280gbmc. The bird data is not publicly available but can be requested from the Dutch Centre for Field Ornithology (Sovon: sovon.nl) or Vigie-Nature (https://www.vigienature.fr/fr/suivi-temporel-desoiseaux-communs-stoc). A summary of the bird data is available via the above Dryad link. The land use maps associated with each scenario have been published in Dellar et al. 2024 (https://doi.org/10.1038/s41597-024-04059-5). For this analysis, all information and results is available via the Dryad repository: https://doi.org/10.5061/dryad.sj3tx96bs.

## Supplementary Material A: Methods

### Choice of WNV host species

The host population for WNV consisted of two groups: high and low competence populations. The ‘high competence’ population was represented by house sparrows, while the ‘low competence’ population was represented by mallards. These species were selected for several reasons. We first selected all bird species which had tested positive for WNV in Europe and restricted this to those i) known to be competent for WNV [64,65]; ii) known to be bitten by *Cx. pipiens* [66]; iii) which have a breeding population in the Netherlands of at least 50,000 [67]; and iv) for which epidemiological parameters were available [68]. Three species remained: mallard (*Anas platyrhynchos*), Eurasian magpie (*Pica pica*) and house sparrow (*Passer domesticus*). We selected mallards and house sparrows for the following reasons: (i) of the three species, magpies have the smallest breeding population in the Netherlands (57,500 vs 230,000 (mallard) and 800,000 (house sparrow) [67]); (ii) mallards and house sparrows show distinct spatial distributions, while the magpie distribution is intermediate (the impact of a uniform distribution is explored in a sensitivity analysis); (iii) there was better species-specific epidemiological data available for mallards and house sparrows than for magpies [68]; (iv) house sparrows and mallards are epidemiologically different with respect to the impact of infection and host competence. Experimental work shows that mallards are less infectious and do not die from infection, while house sparrows are more infectious and can die from infection [36]. In the same study, magpies showed intermediate infectiousness and died from infection.

### Model parameters

Parameter values are described in Table 1 & Table 2. These can be time-dependent (t) or temperature-dependent (T). Most model parameters were directly obtained from literature [68], however some were based on additional calculations.

#### Bird birth rate

The bird death rate was assumed to be constant over space and time, while births were assume to occur seasonally. These births were distributed uniformly across the period where newborns start leaving their nests assuming a hatching period of 14 days and a further 14 days until they fledge the nest [69].

We used a Leslie matrix to calculate the annual number of births assuming the population remains constant in the absence of infection. The following Leslie matrix was developed for the blackbird population, where F stands for the annual fertility rate and S for the annual survival rate of juvenile (j) or adult (a) birds:

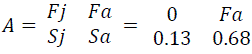

We calculated the fertility rate for adults for which the eigenvalue of the matrix equals 1. The resulting fertility rate was 2.5 births per year, using juvenile survival estimates from [38] and adult survival estimates from [39]. We calculated the daily number of births during the breeding period based on the population size at the beginning of the study period (2.5 births / 45 days = 0.056 births per bird per day).

Annual survival probabilities were transformed to daily mortality rates based on the following equation: *annual survival probability = (1 − daily mortality rates)*^365. Fertility and mortality rates were used to determine the bird population size at each day and location. The same approach was used for all other bird populations.

#### Diapause

The proportion of active adult mosquitoes (ϑ(*t*)) is dependent on daylength φ(*t*):

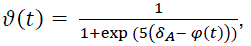

where δ_*A*_ is the autumn threshold when the mosquito population starts entering diapause [70]. While estimates of the autumn threshold value are lacking for the Netherlands, this value was estimated to be 13 hours of daylight in the United Kingdom [70]. Using this threshold with local estimates of daylength suggests that mosquitoes start entering diapause around the beginning of September and all mosquitoes are in diapause towards the end of September. Diapause induction is not only dependent on day length, but also on temperature [61]. To adjust for the somewhat warmer autumn temperatures in the Netherlands compared to the United Kingdom [71,72] and to match observations from local mosquito trapping data, we assumed that the average time to entering diapause (since 1^st^ September) was 20 days, leading to an estimated rate of 0.05 at which mosquitoes enter diapause.

#### Host abundance scaling parameter

Bird and mosquito abundance estimates were represented on a relative scale, because the total population size is difficult to estimate exactly for these populations [73,74]. We therefore introduced a host abundance scaling parameter, which converts the host abundance to be on the same scale as the vector abundance, meaning that their ratio reflects the vector-to-host ratio. This parameter was also used in de Wit et al. [30] and the value was obtained from this study. The value of this parameter is based on the reference scenario of USUV and was applied to WNV following from the assumption that the size of the host population for both diseases is similar. Furthermore, the value of the scaling parameter was kept equal between all scenarios. This allows for changes in bird population size to result in changes in vector-to-host ratios.

#### Estimation of transmission probability house sparrow & mallard to mosquito

Direct estimates of the transmission probability from house sparrow and mallard to mosquito were not available from literature. We estimated these values by combining viremia estimates from experimental infections (Table 2 in [36]) for the two species with the relationship between viremia and transmission probability (Figure 4 in [53]). Kain et al. [53] synthesised literature to estimate the percentage of mosquitoes that became infected following a feeding event on infected blood for different doses. For each day post infection, we extracted the mean West Nile virus viremia levels from [36]. Using the relationship between viremia titre and bird-to-mosquito transmission probability from [53] we determined the transmission probability for each day post infection and then averaged this over the duration of the infectious period.

**Figure 4:**
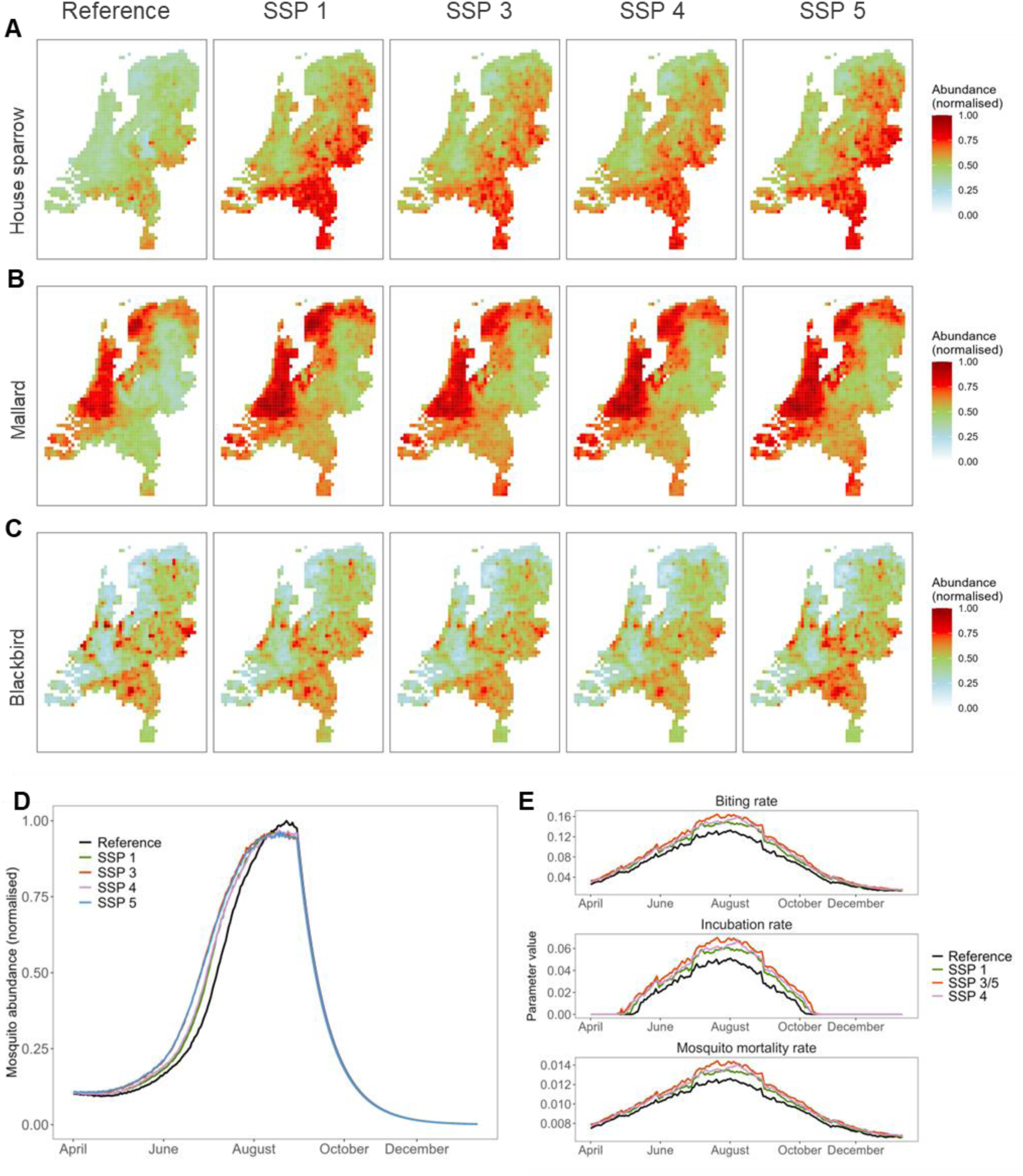
Present and future (year 2050) estimates of breeding season abundance for. (A) House Sparrows, (B) Mallards, and (C) Blackbirds, (D) Cx pipiens abundance, (E) temperature-dependent parameters. Spatial visualizations of mosquito abundance and temperature are available in the supplement.

### Derivation of reproduction numbers

Reproduction numbers were calculated using the Next Generation Matrix (NGM) approach. The reproduction number was obtained by calculating the dominant eigenvalue of the NGM. Below we provide details on the construction of the NGM for USUV. The structure of the NGM for WNV was identical with the only differences being the parameter values and identity of host species. The construction of this NGM has previously been described by de Wit et al. [30], but is repeated here for completeness. The dimensions of the NGM were 5×5 representing the number of species capable of transmitting the infection (mosquitoes (M), juvenile blackbirds (J), adult blackbirds (A), juvenile reservoir population (R), adult reservoir population (P)). Each element in the NGM, *k*_ij_, represents the expected number of cases of type *i* caused by an individual of type *j*:

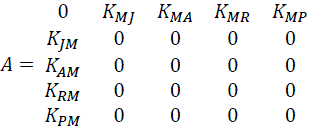

All mosquito-to-mosquito and bird-to-bird elements in the NGM are set to zero as transmission only occurred between mosquitoes and birds and vice versa. The reproduction number is given by 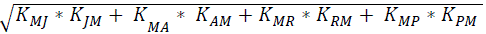. The value of each of the elements in the NGM was derived as follows:

K_JM_: The number of infected juvenile blackbirds caused by an infectious mosquito is determined by

- The fraction of mosquitoes transitioning from E (exposed) to I (infectious)
- The duration of infectiousness in mosquitoes
- The mosquito-to-juvenile blackbird transmission rate

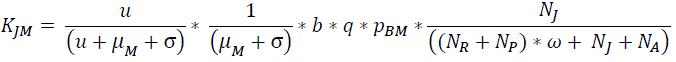

K_AM_: The number of infected adult blackbirds caused by an infectious mosquito is determined by

- The fraction of mosquitoes transitioning from E to I
- The duration of infectiousness in mosquitoes
- The mosquito-to-adult blackbird transmission rate

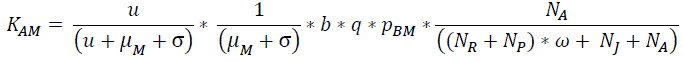

K_RM_: The number of infected juvenile reservoir hosts caused by an infectious mosquito is determined by

- The fraction of mosquitoes transitioning from E to I
- The duration of infectiousness in mosquitoes
- The mosquito-to-juvenile reservoir transmission rate

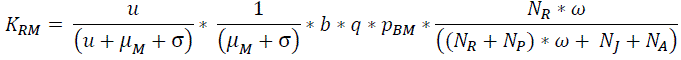

K_PM_: The number of infected adult reservoir hosts caused by an infectious mosquito is determined by

- The fraction of mosquitoes transitioning from E to I
- The duration of infectiousness in mosquitoes
- The mosquito-to-adult reservoir transmission rate

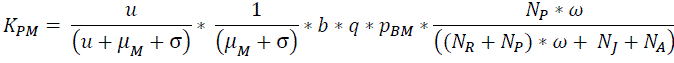

K_MJ_: The number of infected mosquitoes caused by an infectious juvenile blackbird is determined by

- The fraction of juvenile blackbirds transitioning from E to I
- The duration of infectiousness in juvenile blackbirds
- The juvenile blackbird-to-mosquito transmission rate

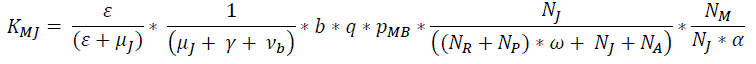

K_MA_: The number of infected mosquitoes caused by an infectious adult blackbird is determined by

- The fraction of adult blackbirds transitioning from E to I
- The duration of infectiousness in adults
- The adult blackbird-to-mosquito transmission rate

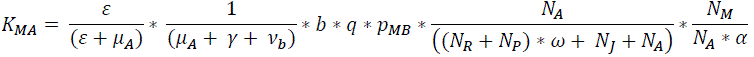

K_MR_: The number of infected mosquitoes caused by an infectious juvenile reservoir host is determined by

- The fraction of juvenile reservoir birds transitioning from E to I
- The duration of infectiousness in juvenile reservoir birds
- The juvenile reservoir-to-mosquito transmission rate

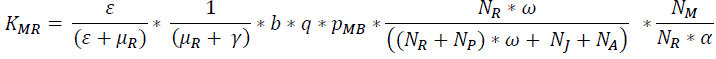

K_MP_: The number of infected mosquitoes caused by an infectious adult reservoir host is determined by

- The fraction of adult reservoir birds transitioning from E to I
- The duration of infectiousness in adult reservoir birds
- The adult reservoir-to-mosquito transmission rate

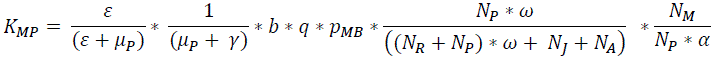

## Supplementary Material B: Additional results

**Figure 1:**
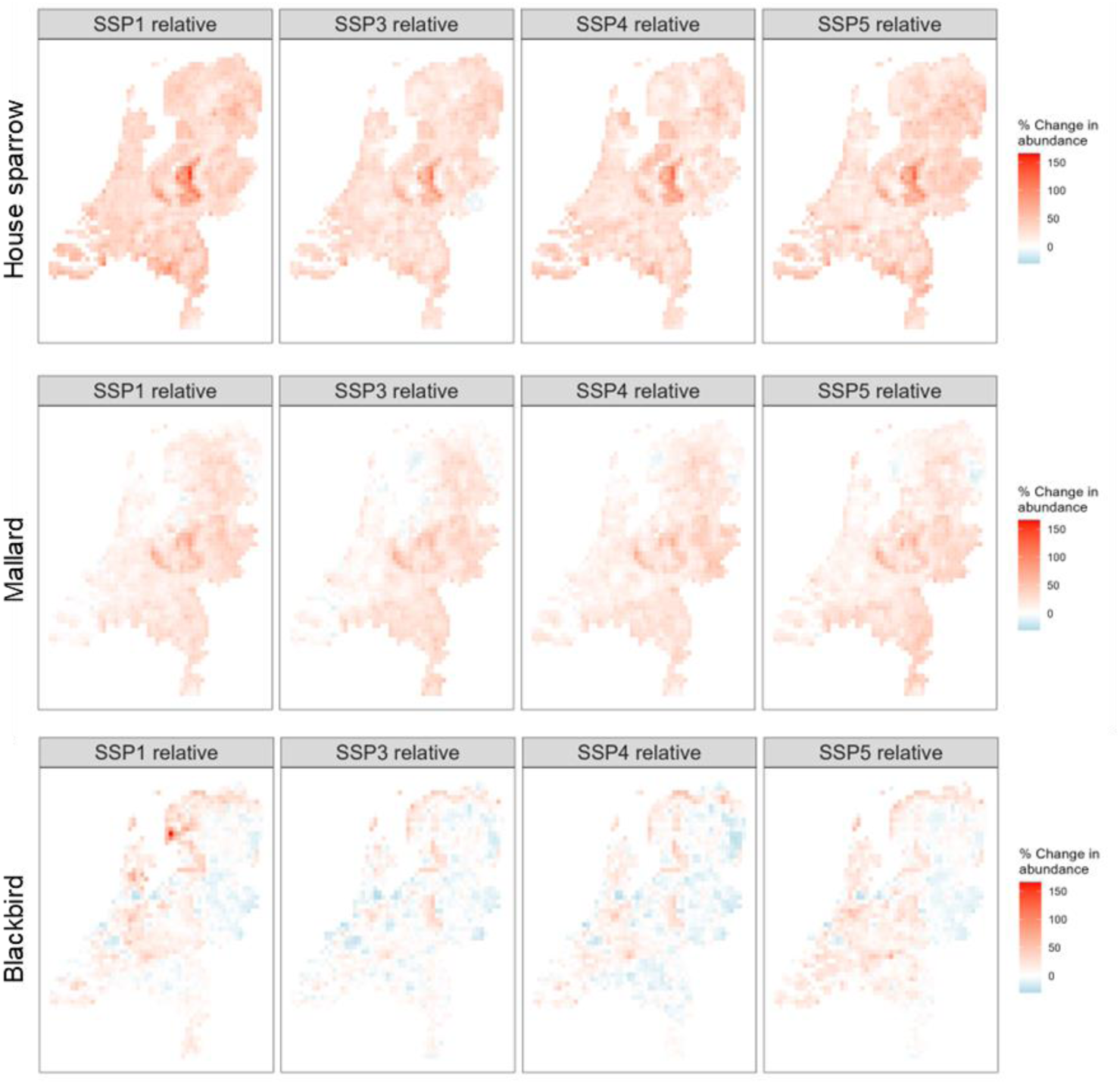
Maps of potential future change (year 2050) in breeding season bird abundance compared to the reference scenario.

**Figure 2:**
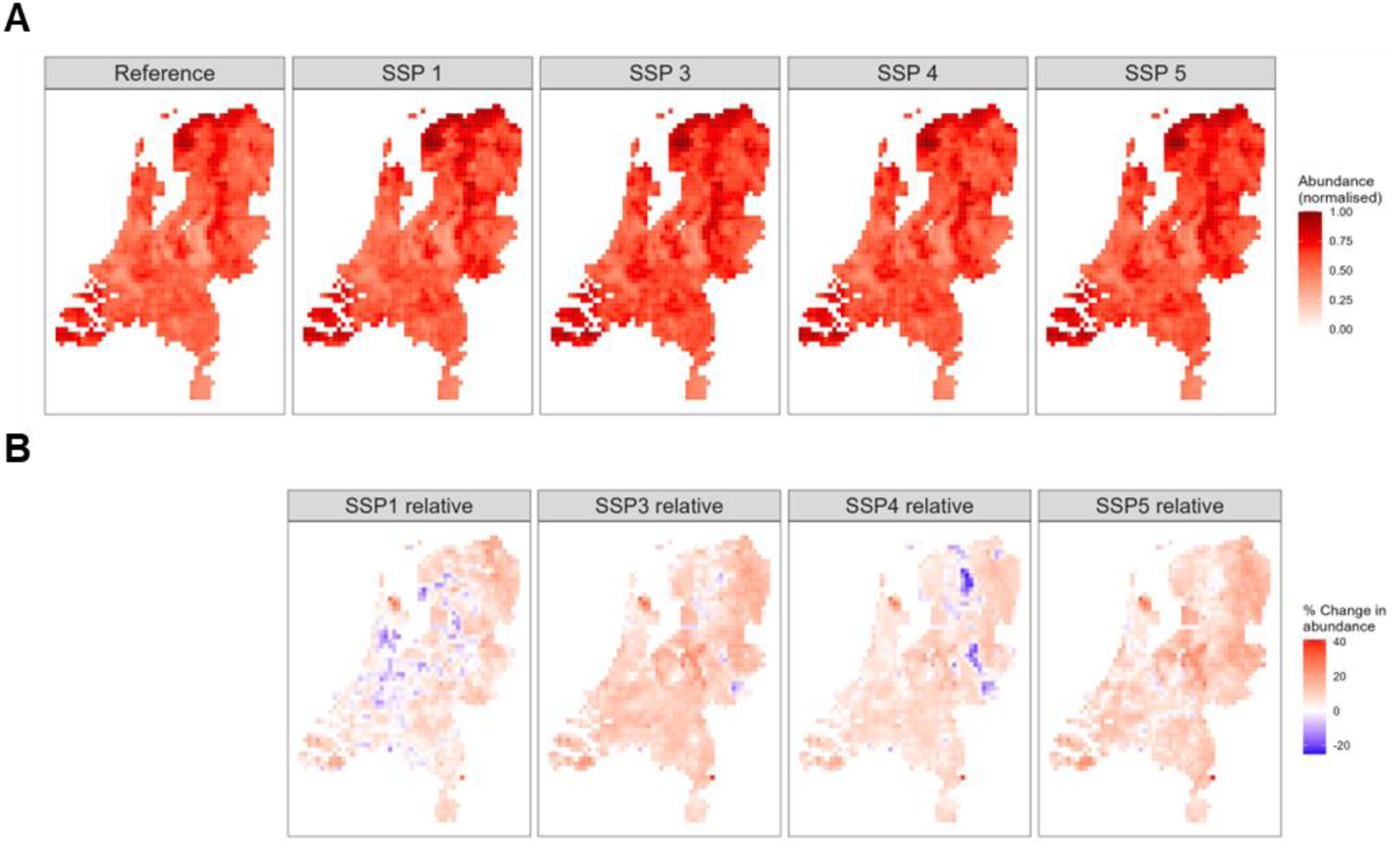
Mosquito abundance per scenario. A) Map of mosquito abundance in reference and future scenarios (year 2050). B) Map of potential future change (year 2050) in mosquito abundance compared to the reference scenario.

**Figure 3:**
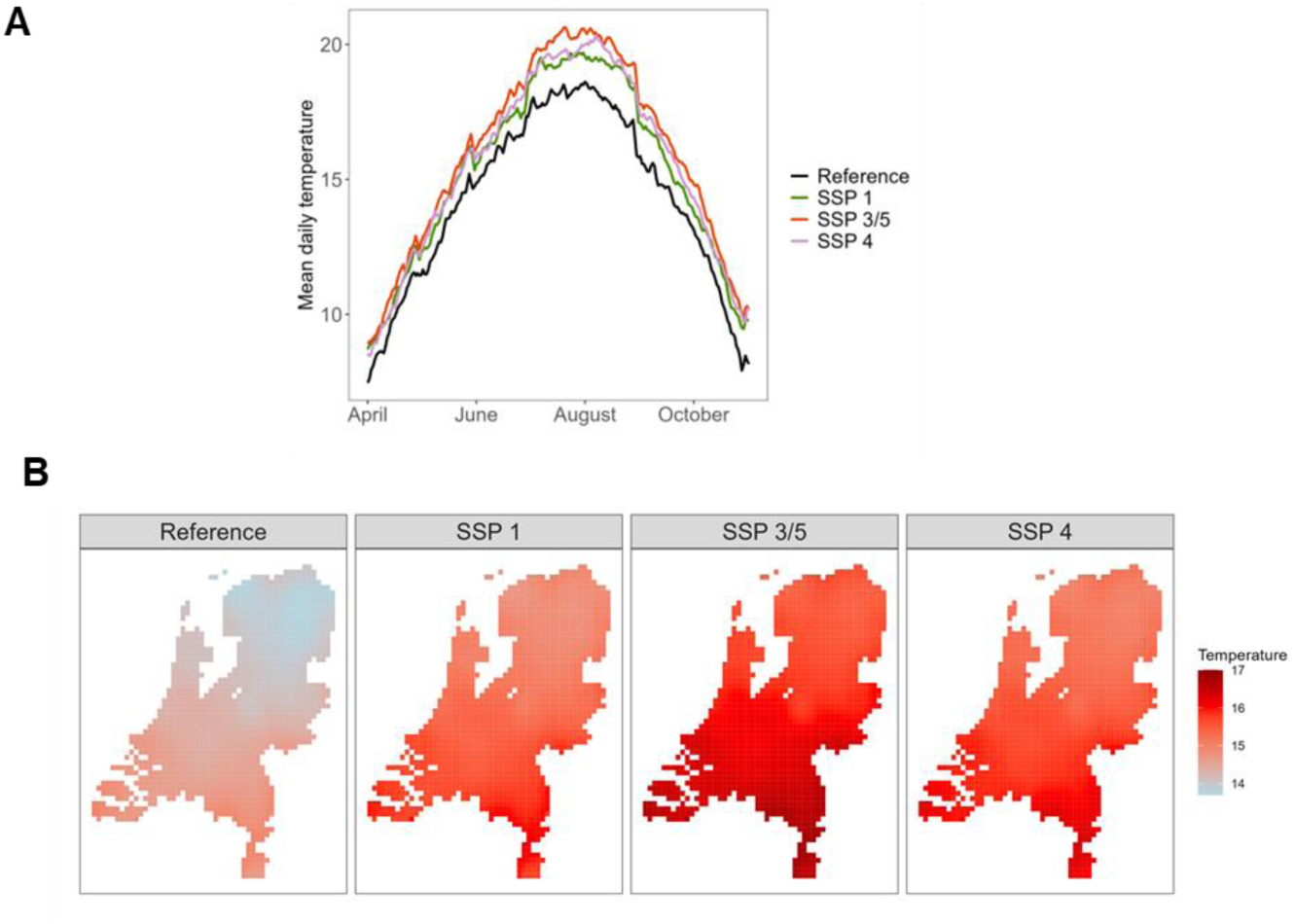
Temperature per scenario. A) National mean daily temperature for reference and future scenarios (year 2050). B) Map of mean daily temperature across April-November for each location for the reference and future scenarios (year 2050).

**Figure 4:**
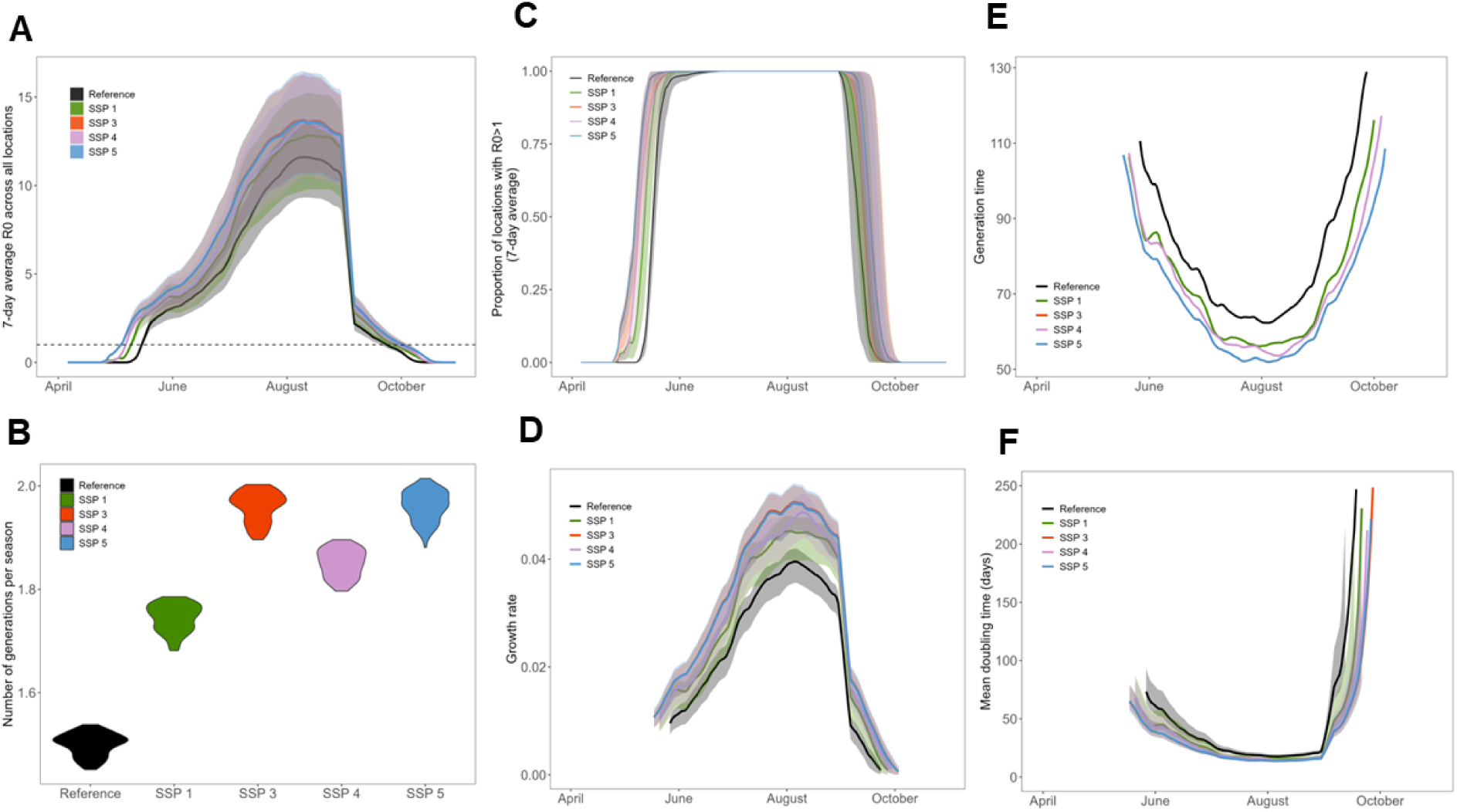
Comparison between reference and all future scenarios for USUV for. (A) R0 over time, (B) the number of generations per season, (C) the proportion of locations with R0>1, (D) the epidemic growth rate, (E) the generation time, and (F) the epidemic doubling time.

**Figure 5:**
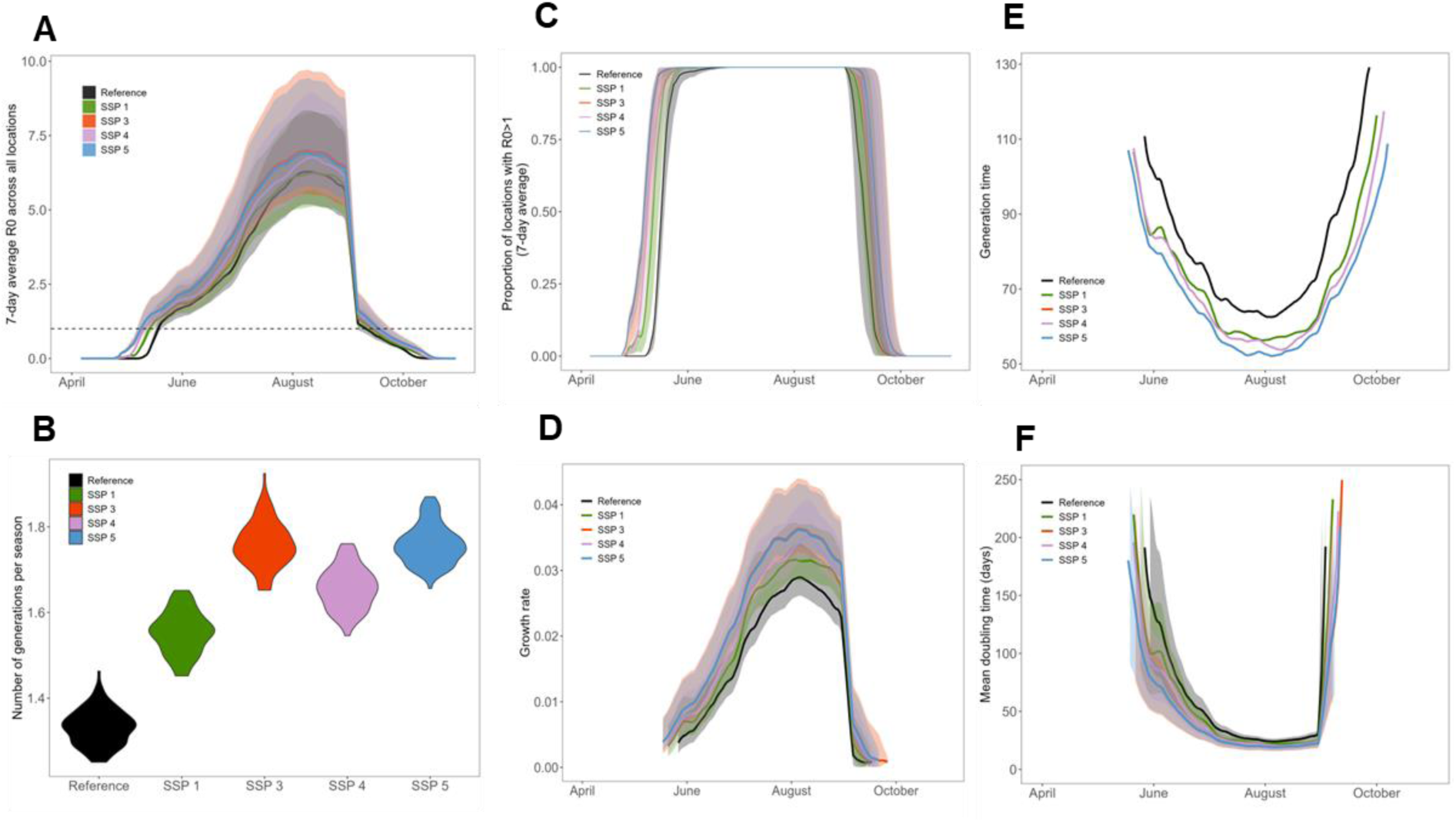
Comparison between reference and all future scenarios for WNV for. (A) R0 over time, (B) the number of generations per season, (C) the proportion of locations with R0>1, (D) the epidemic growth rate, (E) the generation time, and (F) the epidemic doubling time.

**Figure 6:**
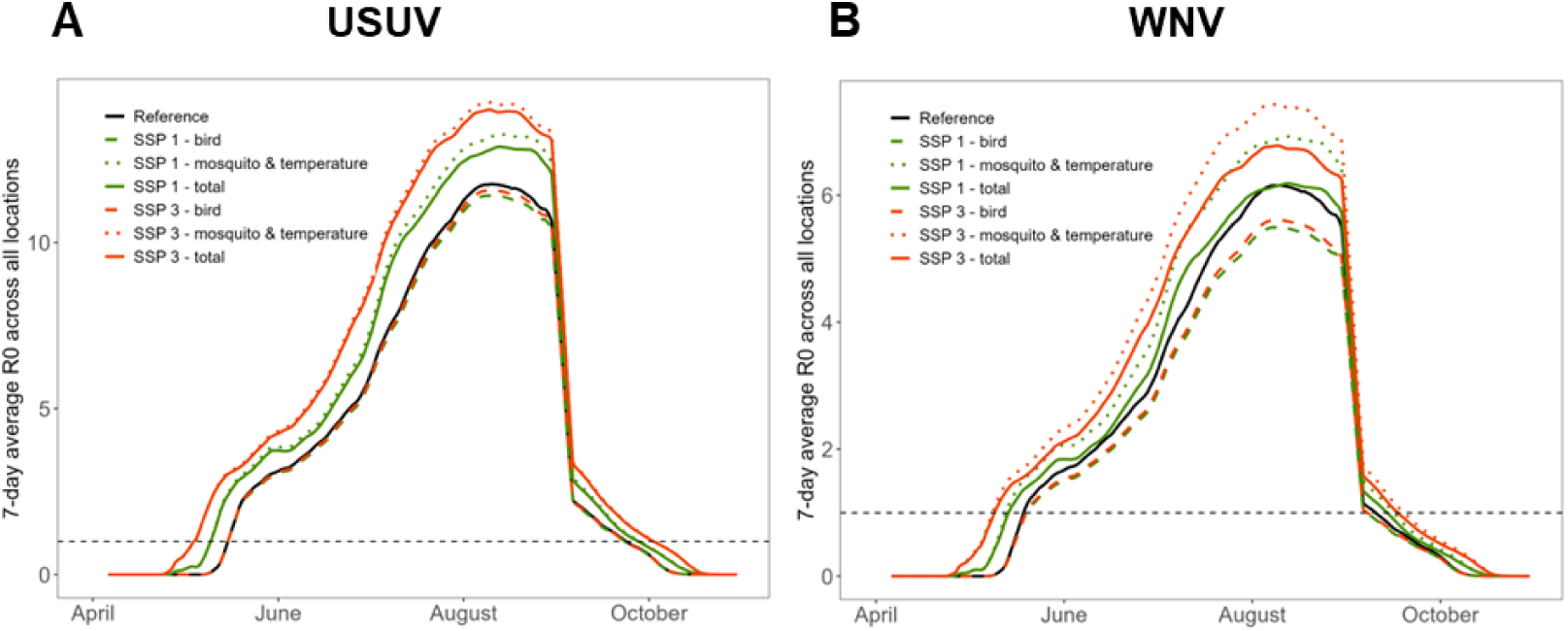
Comparison of R0 over time between reference and split-scenarios for USUV. (A) and WNV (B). ‘Bird’ scenarios represent scenarios where only the bird abundance changes, while temperature and mosquito abundance remained equal to the reference scenario. ‘Mosquito & temperature’ scenarios represent scenarios where only the temperature and mosquito abundance change. ‘Total’ scenarios represent the standard scenarios where are elements change.

**Figure 7:**
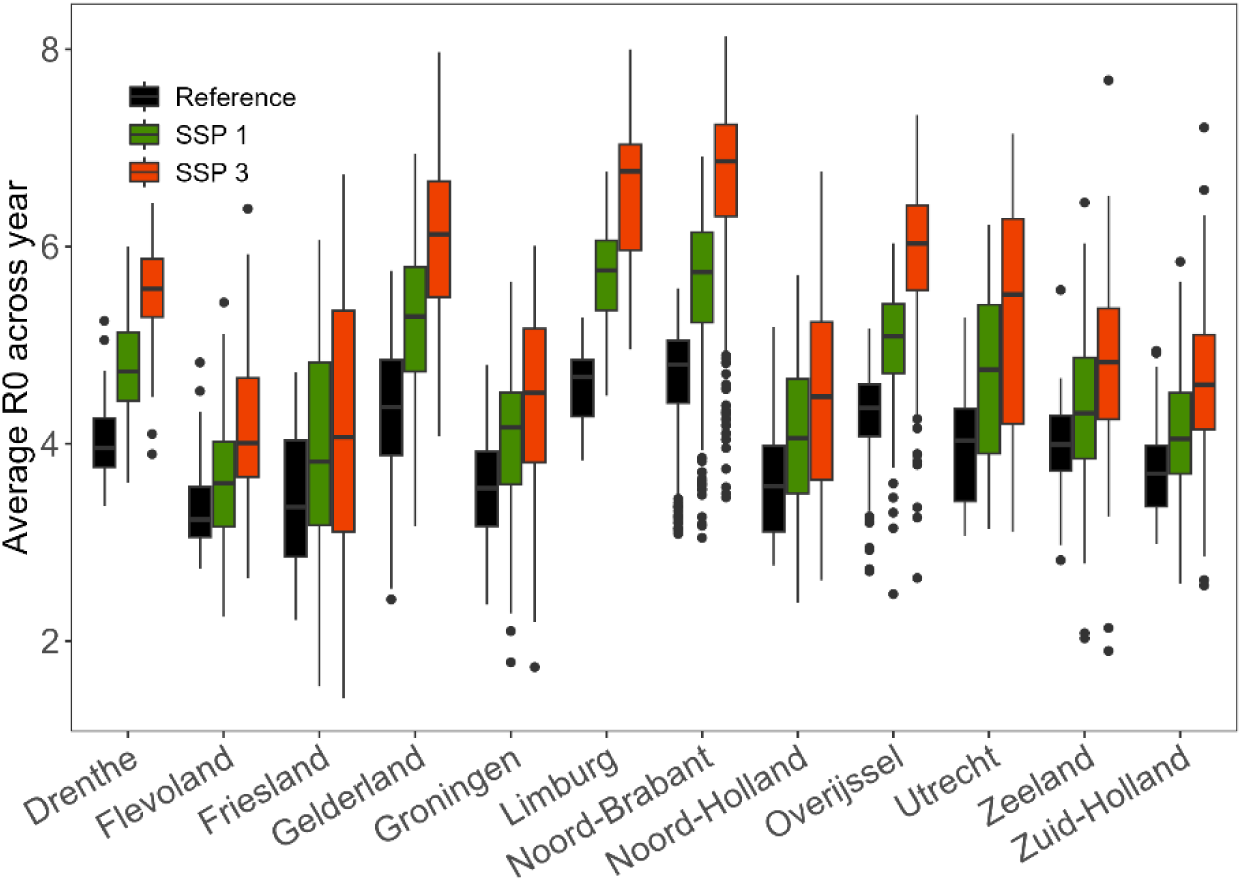
USUV R0 averaged across the year per province for each main SSP scenario.

**Figure 8:**
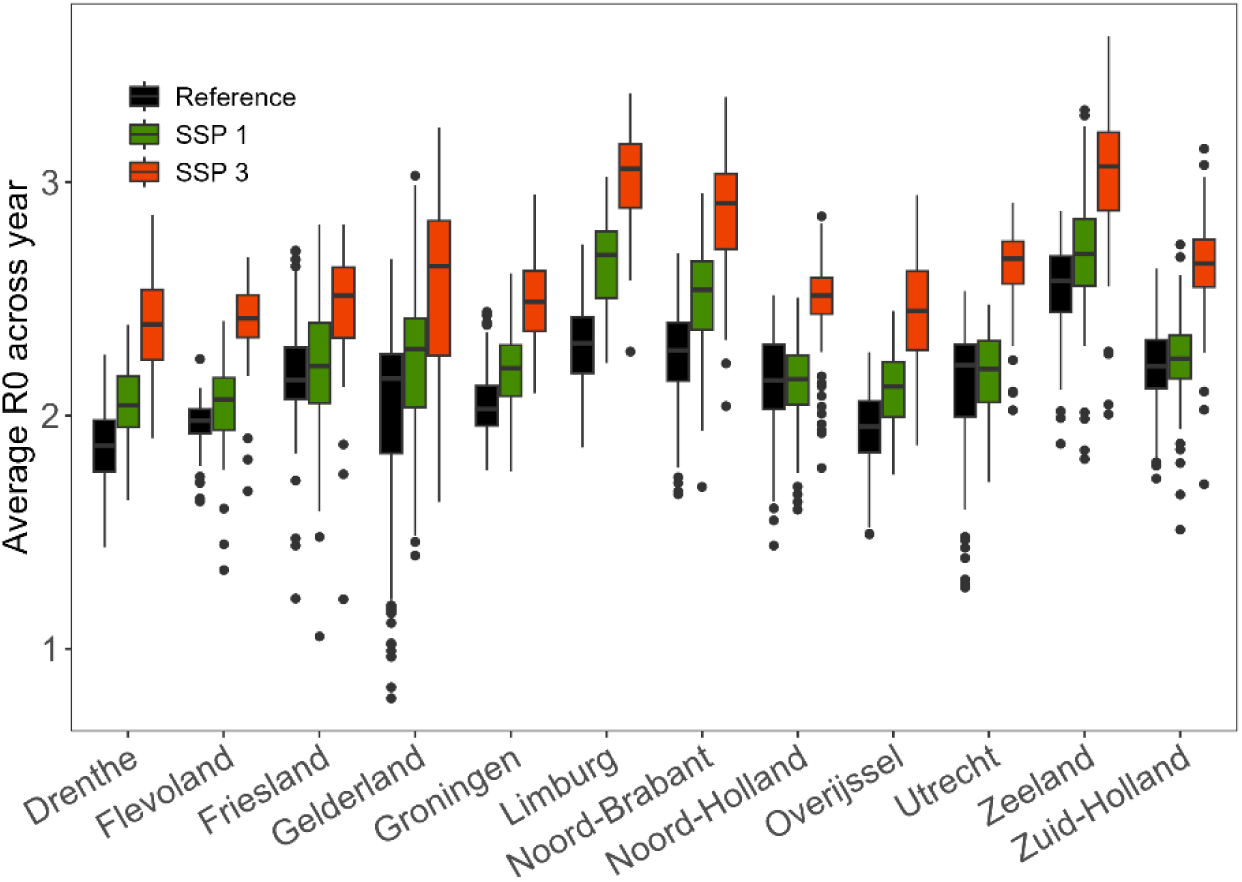
WNV R0 averaged across the year per province for each main SSP scenario.

**Figure 9:**
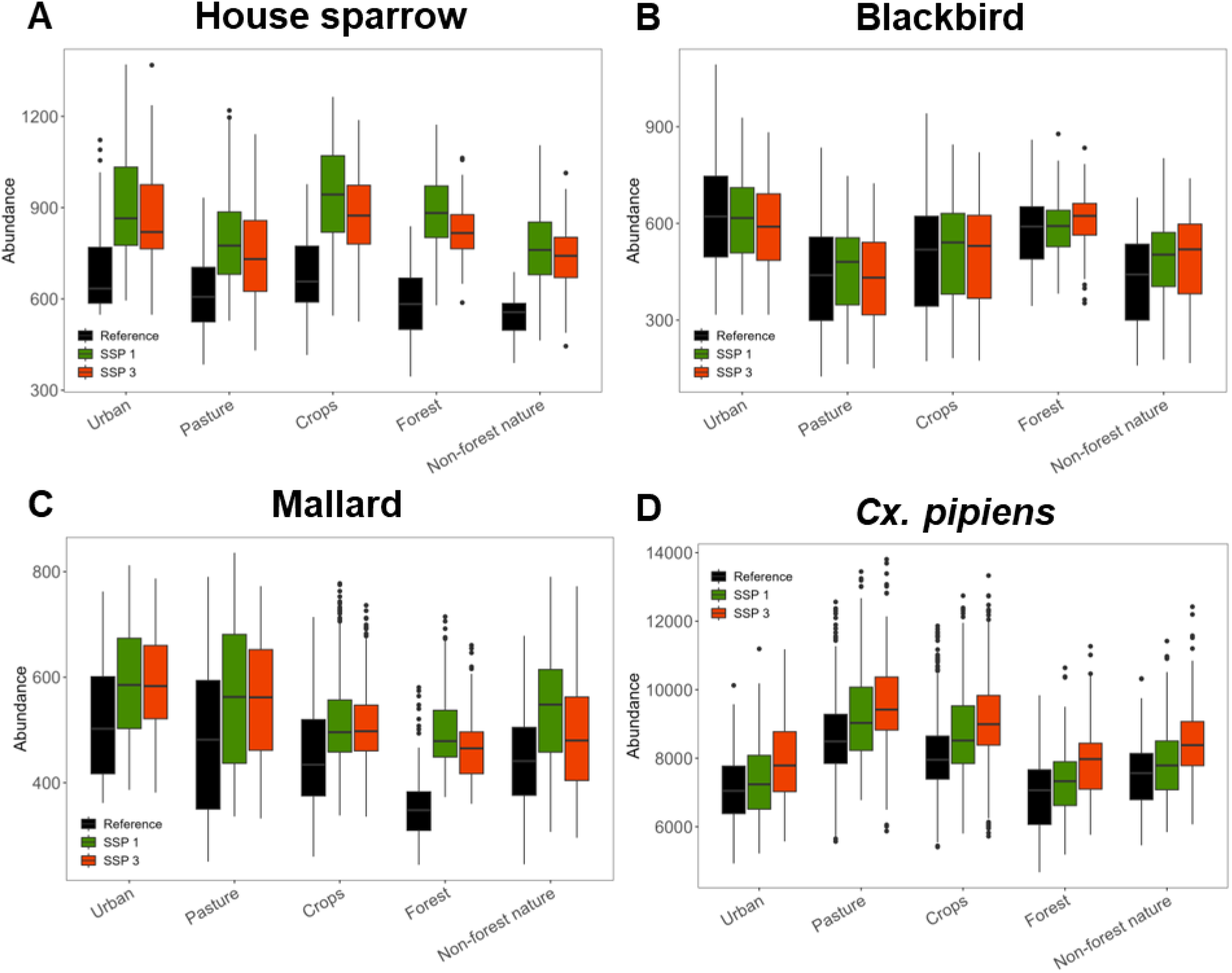
Bird (A-C) and mosquito (D) abundance per land use class for each main SSP scenario.

**Figure 10:**
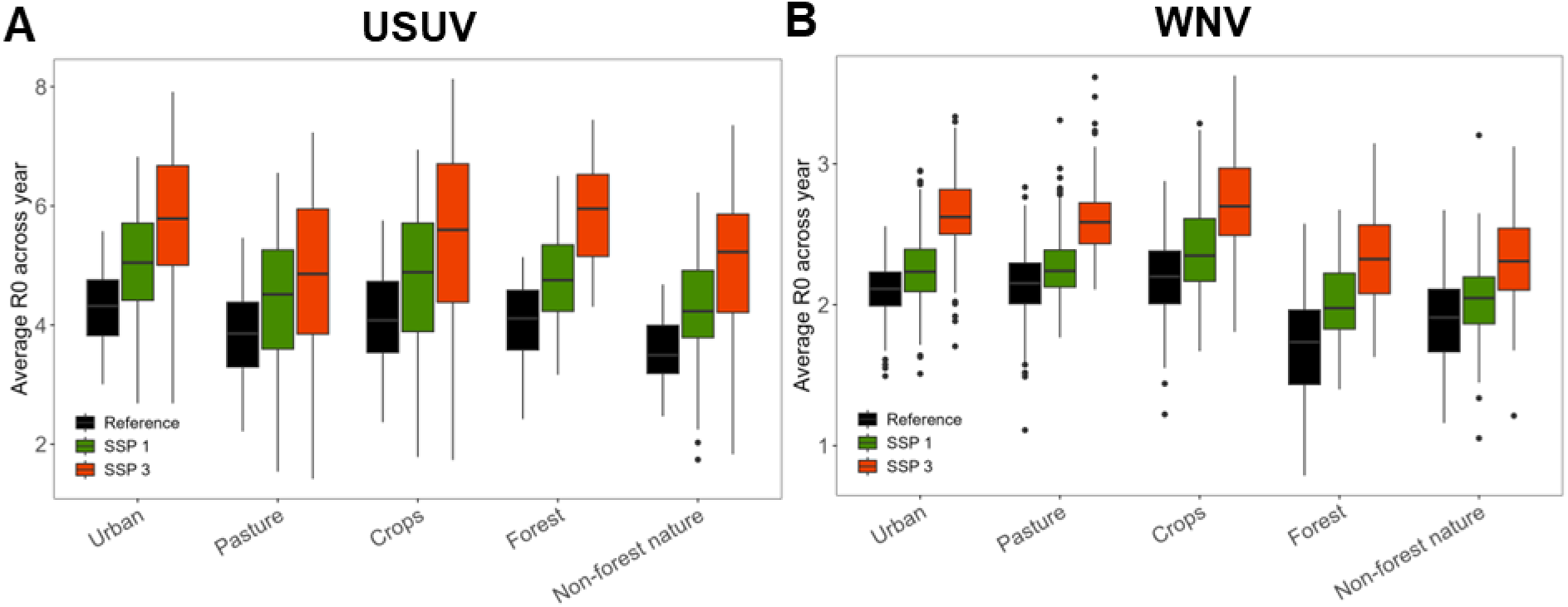
R0 averaged across the year per land use class for each main SSP scenario for (A) USUV and (B) WNV.

**Figure 11:**
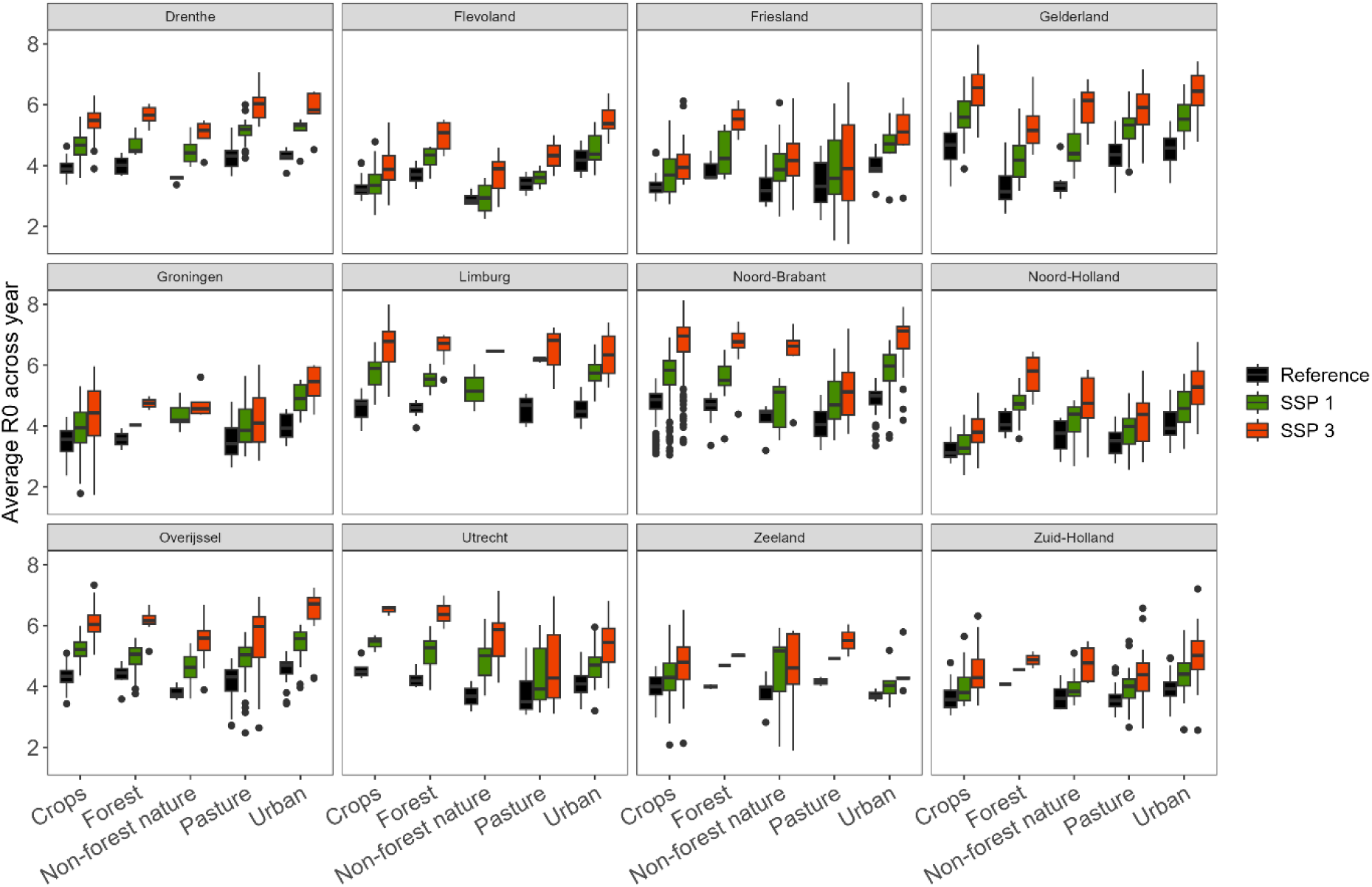
USUV R0 averaged across the year per land use class & province for each main SSP scenario.

**Figure 12:**
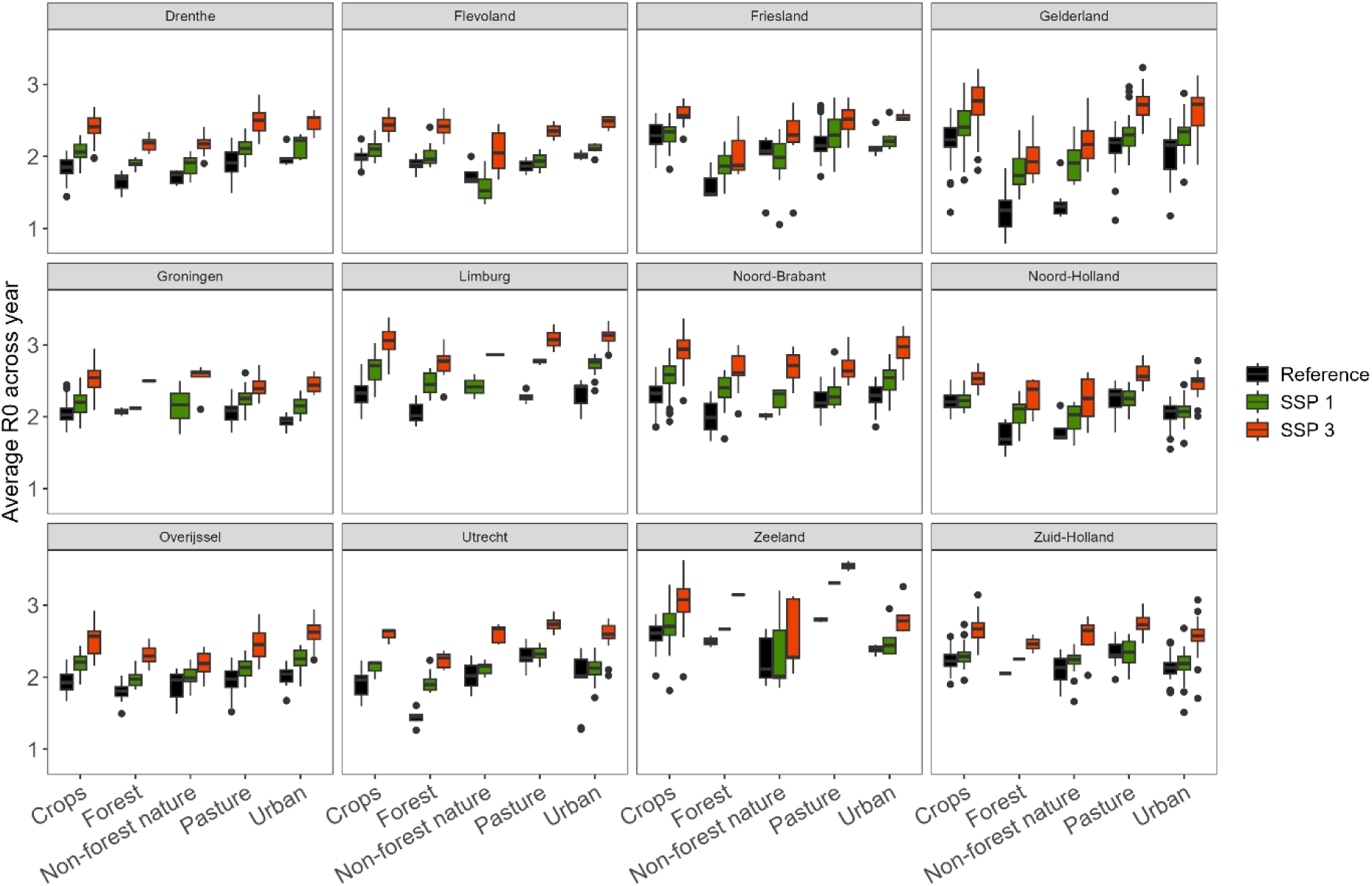
WNV R0 averaged across the year per land use class & province for each main SSP scenario.

## Supplementary Material C: Sensitivity analyses

**Table 1:**
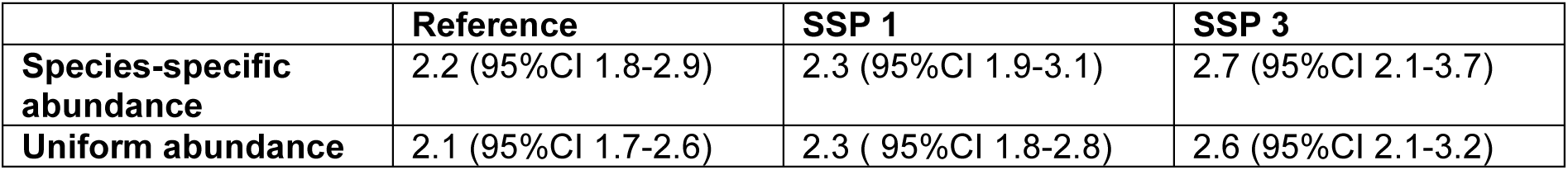
Comparison of average R0 across (all locations and days) between using the species-specific abundance distribution and a uniform bird distribution.

**Figure 13:**
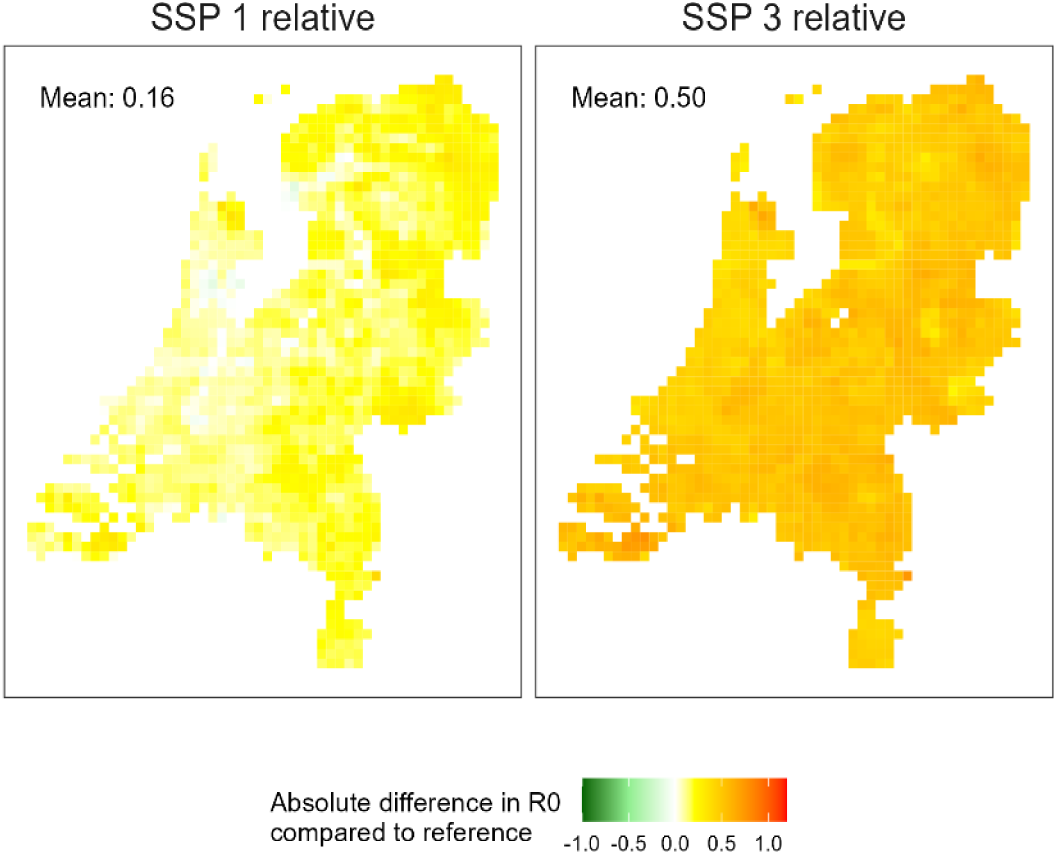
Map of the difference in WNV basic reproduction number per grid cell between scenarios and the reference. WNV hosts were uniformly distributed across the country in this analysis.

In this sensitivity analyses we explored the impact of assuming a uniform spatial distribution of WNV hosts. While the highest R0 values in the reference scenario were still observed in Zeeland, the increases in R0 values in future scenarios were less heterogeneous with relative increases ranging from 1.03 times in Noord-Holland to 1.12 times in Groningen for SSP1 (1.20 and 1.26 for SSP3, respectively) (supplementary Figure 14). Increases in R0 by land use class were also less heterogeneous and ranged from 1.07 in urban and non-forest nature to 1.09 in forest in SSP1 (1.23 in non-forest nature to 1.27 in forest for SSP3) (supplementary Figure 15).

**Figure 14:**
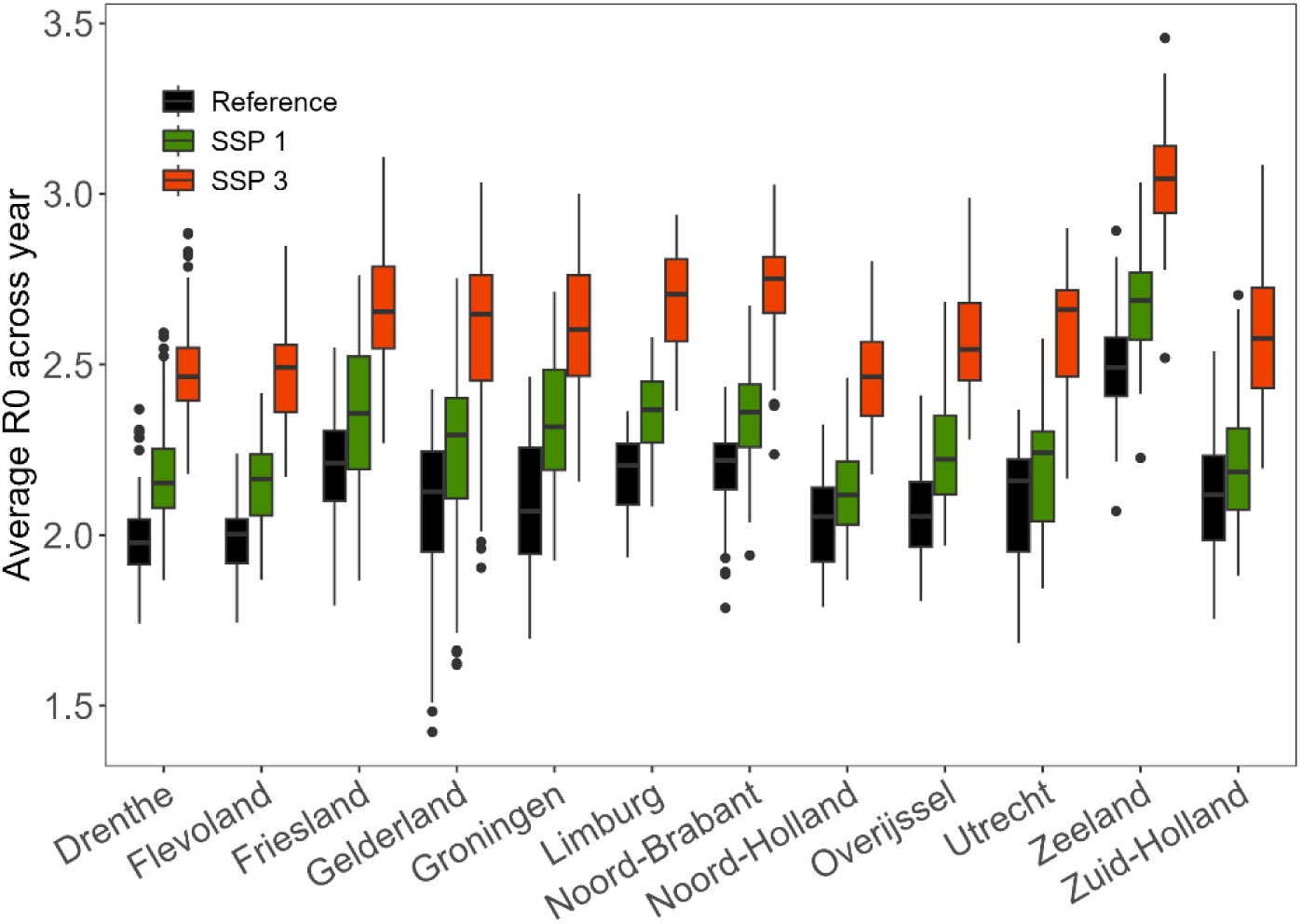
WNV R0 averaged across the year per province for each main SSP scenario. WNV hosts were uniformly distributed across the country in this analysis.

**Figure 15:**
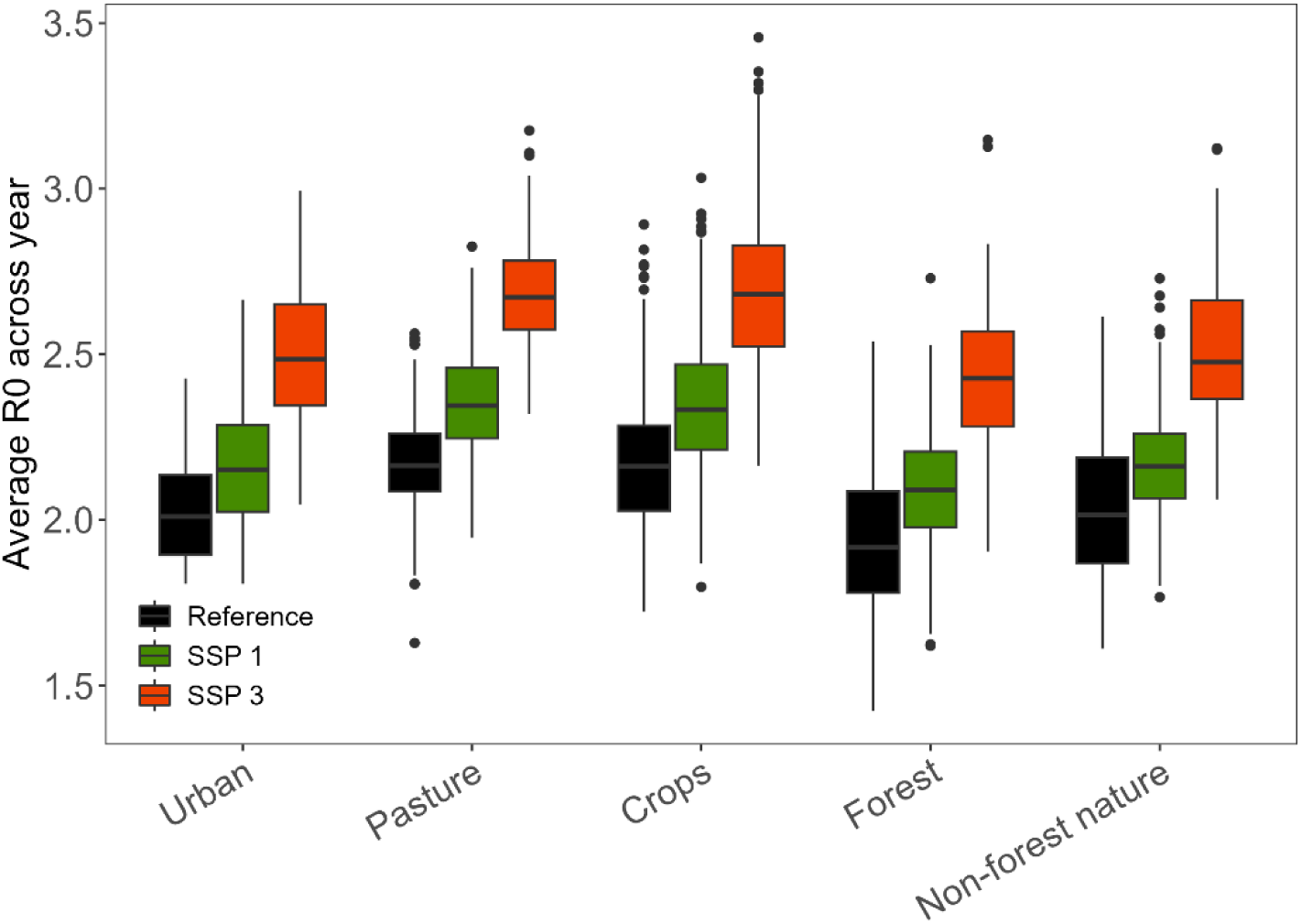
R0 averaged across the year per land use class for each main SSP scenario for WNV. WNV hosts were uniformly distributed across the country in this analysis.

